# Mapping cellular nanoscale viscoelasticity and relaxation times relevant to growth of living Arabidopsis thaliana plants using multifrequency AFM

**DOI:** 10.1101/2020.06.30.180463

**Authors:** Jacob Seifert, Charlotte Kirchhelle, Ian Moore, Sonia Contera

## Abstract

The shapes of living organisms are formed and maintained by precise control in time and space of growth, which is achieved by dynamically fine-tuning the mechanical (viscous and elastic) properties of their hierarchically built structures from the nanometer up. Most organisms on Earth including plants grow by yield (under pressure) of cell walls (bio-polymeric matrices equivalent to extracellular matrix in animal tissues) whose underlying nanoscale viscoelastic properties remain unknown. Multifrequency atomic force microscopy (AFM) techniques exist that are able to map properties to a small subgroup of linear viscoelastic materials (those obeying the Kelvin-Voigt model), but are not applicable to growing materials, and hence are of limited interest to most biological situations. Here, we extend existing dynamic AFM methods to image linear viscoelastic behavior in general, and relaxation times of cells of multicellular organisms in vivo with nanoscale resolution, featuring a simple method to test the validity of the mechanical model used to interpret the data. We use this technique to image cells at the surface of living *Arabidopsis thaliana* hypocotyls to obtain topographical maps of storage E’ = 120 − 200 MPa and loss E’’= 46 − 111 MPa moduli as well as relaxation times τ = 2.2 − 2.7 µs of their cell walls. Our results demonstrate that cell walls, despite their complex molecular composition, display a striking continuity of simple, linear, viscoelastic behavior across scales–following almost perfectly the standard linear solid model–with characteristic nanometer scale patterns of relaxation times, elasticity and viscosity, whose values correlate linearly with the speed of macroscopic growth. We show that the time-scales probed by dynamic AFM experiments (milliseconds) are key to understand macroscopic scale dynamics (e.g. growth) as predicted by physics of polymer dynamics.

## 1 Introduction

Biology constructs complex hierarchical structures through coordination of biochemical and mechanical factors across multiple spatio-temporal scales during morphogenesis. From the standpoint of physics, biological dynamic shapes result from life’s use of energy: organisms create and preserve structures without violating the second law of thermodynamics by dissipating energy into the environment (e.g. as heat) [1, 2]. In mechanical terms this means that biological structures are not elastic–they do not respond instantaneously to perturbations like ideal Hookean springs. They are viscoelastic, i.e. they react to forces with characteristic time-scales, which emerge from their energy dissipation patterns at the nanoscale.

Recently developed dynamic AFM imaging methods are capable of mapping viscoelasticity of living biological systems and organic materials with nm resolution [3, 4]. However, there exists a problem for AFM quantitative mapping techniques in their current form: they utilize the Kelvin-Voigt (KV) linear viscoelasticity model (a Hookean spring connected in parallel to a Newtonian viscosity dash-pot) to interpret AFM observables. In materials following the KV viscoelasticity model, deformations are always reversible - hence the KV model is not appropriate to describe growing materials, and inapplicable to a large proportion biologically relevant situations. Additionally, existing quantitative AFM imaging techniques lack methods to test if the viscoelastic model utilized is valid for a particular material, and therefore their usefulness remains limited, especially in biological contexts. Without a test of the validity of the model used to interpret the data, the value of the results remains questionable, and this is one of the main challenges that we address in this paper.

Here, we aim to exploit the frequency range of dynamic AFM and develop an AFM technique that is able to correctly map the mechanical properties of a biological material at the nm and cellular scales. To achieve it we expand an existing multifrequency AFM technique [5] so that we are able to simultaneously map topography and time-relaxation in living eukaryotic cells with nm-precision; we additionally provide a novel method to test the validity of the viscoelastic model used to extract quantitative maps of relevant quantities. Our technique is also able to non-invasively map energies stored/dissipated, viscosity, and elasticity of living cells in their tissue context. We use *Arabidopsis thaliana* plants as model systems. From the biological physics standpoint, plants are ideal systems for investigating the mechanics underlying morphogenesis *in vivo*, as growth is dominated by the mechanics of the cell wall (CW) surrounding each cell. Plants continuously grow and shape new organs by regulating cell proliferation, growth rate, and growth direction over time. Growth is driven by the cells’ undirected internal turgor pressure, but molecular modification of the CW is required to allow directional growth while maintaining structural integrity and withstanding turgor pressure [6-10]. As a consequence of their central role in growth control, the biochemical and mechanical properties of plant CWs are of great interest to the plant scientific community and have been extensively investigated. From a biochemical perspective, it has long been known that the CW comprises a largely of carbohydrates (cellulose microfibrils, pectins, hemicelluloses) and a small fraction of structural proteins [11], although our understanding of the sophisticated fibre-reinforced network formed by these polymers is still in flux and has recently undergone major revisions [12]. However, it is clear that CW’s mechanical properties emerge from the biomechanical and biochemical properties of these individual components and their interactions, which are modulated by hormonal/enzymatic activity and active changes of structural anisotropy [12-16]. Changes in cell wall mechanics have been linked to morphogenesis in many studies [14, 17, 18], reviewed in [19], although the precise relationship between cell wall mechanical properties and growth is not well understood [12]. One reason for this is that the different techniques used to probe cell wall mechanical properties (including tensile stretching of whole tissue or excised cell walls [20][21], optical techniques like Brillouin light-scattering microscopy [22, 23] and indendation-based techniques like AFM) measure different quantities and produce widely differing results [12, 19]. In particular, CW elastic moduli obtained using indentation techniques like AFM and cellular force microscopy (CFM) [13, 14] in response to growth factors, turgor pressure, pectin methyl esterification, or enzymatic activities [14-18, 24, 25] are typically orders of magnitude lower than those obtained in tensile stretching experiments or osmotic pressure measurements [12, 19], raising the question of how these measurements relate to growth.

Furthermore, with a few exceptions [26], AFM studies on plant CWs have largely been quasi-static, and thus limited to extracting elastic moduli, and the measurement of time dependent mechanical properties has remained largely out of experimental reach at the subcellular level. However, CWs are viscoelastic, and the time-dependent component of its mechanical properties created by molecular structures across time and length-scales is of critical interest in the context of growth. Although the whole plant growth rates are in the order of seconds/minutes, the relevant time scales for the growth of the cell wall are expected to be distributed in a wide, continuum spectrum. The CW mechanical behavior emerges from interactions between polymer chains and molecules that form the CW and are characterized by a large range of different time and length scales that depend, for example, on the mechanism of action of the enzymes [27]. Real-time imaging has revealed that cellulose microfibers reorient in the second/minute scale, probably as a result of the effect of turgor pressure-induced wall elongation, and facilitated by the activity of wall-loosening enzymes [28], the resolution of these studies did not allow to image dynamics at lower time/space scales. Apart from the biochemistry underlying e.g. the loosening of the cell wall [13], there are relevant mechanical properties that emerge from the physics of the CW polymeric composite, that have not been probed at the time scales that characterize their dynamics. Polymer physics theory has been particularly successful at relating the structure and connectivity of the polymer chains in a polymeric material with the time-scales that are relevant to explain the mechanical behavior at larger scales [27, 29]. In experimental polymer physics, a spectrum of time-scales (1-100s Hz) is probed by spectroscopic techniques such as dynamic mechanical analysis (DMA)[29], this allows to connect the properties of a macroscopic sample with the molecular dynamics of the polymer, however macroscopic DMA is not applicable to plant CWs. In particular, polymer physics has shown that the full polymer chain length (characterized by the radius of gyration Rg) is related to polymer dynamics time scales, in the particular case of the polymers in the plant CW Rg = 10 – 100 µm, which means their associated time-scales are of the order milliseconds[27]. The millisecond time scale should be relevant at the nanometer scale, while the larger rearrangements of cell wall material previously measured would happen in larger times scales. One of the characteristics of polymeric materials that makes them so suitable to construct biological shapes is that they are able to connect molecular short time scales with the slower dynamics of the large structures that they form [27]. There are few techniques that can be used to estimate the mobility of the polymers within the CW at sub-second timescales. Solid State NMR spectroscopy has been used to study the dynamics of the CW polysaccharides using the spin-lattice relaxation of ^13^C. This method is based on the local motion of the ^13^ C nucleus and can give an indication for the relative average mobility of a specific macromolecule, but has no spatial resolution and the link between the relaxation times of the nucleus of ^13^C and the overall relaxation times of the dynamics of macromolecules is not established [30, 31]. Recently dynamic AFM techniques have been able to probe the mechanical properties of living systems in the kHz (millisecond) range. These techniques usually indent the sample with a nanometer size tip and over-impose one or more frequencies on the indentation. Using suitable theories, it is possible to extract time-dependent mechanical properties of the sample by using the AFM cantilever observables (amplitude, indentation, phase, etc) [4]. Advanced quantitative mechanical property mapping multifrequency AFM techniques have been used for mapping of local viscoelastic properties of living cells at high imaging speeds[4, 5, 32] and kHz frequencies, although they have not been applied to living tissues. One of the main limitations of current dynamic AFM measurements is the unmet requirement to incorporate both the correct theoretical model to extract mechanical properties from the AFM observables (Suplementary Information SI, S1) and a suitable test of the validity of the model utilized. In the case of CWs, the model used to interpret must describe mechanical characteristics of a growing material (i.e. the Maxwell (MW) model, the more general standard-linear-solid model, SLS, or the generalized MW model [10, 26, 33-35]. The research done so far has not addressed this fundamental issue.

Here, we present a simple theoretical framework that overcomes existing experimental difficulties with existing AFM measurements including quantitative agreement with macroscopic values of elasticity. We apply our quantitative AFM mechanical property imaging method to cells present at the surface of the hypocotyl (an embryonic organ connecting root and shoot) of Arabidopsis thaliana. We have chosen this organ of the plant because cell division rarely occurs in it, therefore, our nanoscale AFM mapping of energy stored/dissipated, viscosity, elasticity, and time relaxation probes only the mechanical properties that underlie pure growth (excluding division).

## 2. Materials and Methods

### 2.1 Atomic force microscopy

All experiments were performed with the Cypher ES (Asylum Research, Santa Barbara, CA) in pure water. The AFM was operated in contact resonance, i.e. the feedback was on the deflection of the cantilever is simultaneously oscillated at the first eigenmode using photothermal actuation. The cantilever used was from Nanosensors (PPP-NCLAuD) with a spring constant of k ≈ 36(3) N m−1 and a resonance frequency in water f ≈ 78(4) kHz and a quality factor (Q) Q = 9.7. The cantilever was calibrated using the Sader method[36]. The scan rate was 2.44 lines/s with 255 pixels/line which corresponds to a pixel size of 78 nm. The free amplitude was set to A1,far ≈ 14 nm with a blue laser power of P_blue_ = 8 mW and the amplitude in contact was about A_1_ ≈ 4 nm with a set point of the deflection of 0.3 V, which corresponds to an indentation depth of about 300 nm. The exact drive frequency was re-tuned before each scan and at the same time the phase far from the surface was set to φ_1,far_ = 90(1). At the end of each scan, a quasi-static indentation curve was obtained in the centre of the image to obtain a calibration curve for A_1,near_, φ_1,near_, and E_0_. A detailed flowchart describing the experimental setup and the use of theoretical models can be found in Fig. S5 (SI).

### 2.2 Plant material and growth conditions

All plants used were Arabidopsis thaliana ecotype Columbia (Col-0). AFM experiments were performed on wild-type plants. For measurements of cellular growth rates, plants expressing the plasma membrane marker YFP:NPSN12[37][36] were used. Seeds were surface sterilised with 70% ethanol and grown on vertically oriented plates containing 2.2 g L^-1^ Murashige and Skoog growth medium (Sigma-Aldrich, pH5.7) supplemented with 1 % sucrose and 0.8 % Bacto agar (BD Biosciences). To assure synchronized germination, seeds were stratified for three days at 4 °C before transfer to a growth chamber (20 °C). Plates were exposed for 6 h to 110 molm-2s-1 white LED light. Plates were then wrapped in a double layer of aluminium foil and kept for 60 h for AFM experiments, or 48 h, 60 h, and 72 h, respectively for cell length measurements. Growth experiments were also presence of 3 nN isoxaben (IXB) in the growth medium; IXB was dissolved in dimethyl sulfoxide (DMSO) before adding to the medium. Growth medium containing the same concentration of DMSO as that present in the IXB experiments was used for control experiments.

### 2.3 Confocal microscopy and image analysis

Confocal images of hypocotyls expressing YFP:NPSN12 were acquired after 48 h, 60 h, and 72 h or at 58h and 60 hours respectively, using a HCX PL APO CS 20/0.7 IMM UV lens on a Leica TCS SP5 confocal microscope using 514 nm excitation and a 525 nm-580 nm emission range (Fig. S10). Image analysis and processing was performed in Fiji [38]: Maximum intensity projections of 3D confocal stacks acquired at consecutive regions along a hypocotyl were assembled using the MosaicJ plugin. Cell lengths were measured from 7-12 cell files from 2-4 hypocotyls for each time point using the lines tool. Average cell length was calculated for four regions (n > 61 cells per region): region 1 (cell 3-5), region 2 (cell 6-8), region 3 (cell 9-11) and region 4 (cell 12-15), and average normalized growth velocity for each region was calculated.

### 2.4 AFM sample preparation

The seedlings were attached to the probe holder of the Cypher, which are 15 mm diameter metal plates, with Hollister 7730 medical adhesive spray. The liquid adhesive was distributed evenly to a thin layer on the plate. The seedling was gently picked up with metal tweezers under the apical hook and placed onto the adhesive. For setting the glue, the attached seedling was placed inside a Petri dish covered in aluminium foil and filled with a wet tissue to keep the plant hydrated. After 15 min a drop of about 50 µl − 100 µl of pure water was placed on the sample, for AFM imaging.

### 2.5 Data Analysis

The AFM data was analyzed entirely with Python3.5 (https://www.python.org/) for which we wrote a library to read the Igor binary wave (ibw) format and process the AFM data with the theory published in this article. The library has been made publicly available at https://github.com/jcbs/ForceMetric.

For the determination of the CWs the topography of the cells was used. The longitudinal walls were then determined by using the fact that the slope of the topography changes signs between adjacent cells. This was implemented by calculating the angle of the slope from the gradient of the topography and applying a canny edge filter to the result. To get the different longitudinal walls regions of interest (ROIs) were defined. This approach was not possible for the transverse walls due to the different topographical properties and therefore, these walls were determined by hand with lines of three or four nodes. Finally, the periclinal walls were determined by the remaining pixels with a margin of 20 pixels around the anticlinal walls. Walls with less than 15 pixels were not considered. Moreover, data which was acquired within 5 % of the limits of the z-piezo were not considered in the analysis to reduce imaging artifacts. Statistical analysis between the different walls was done with a two-tailed Welch’s t-test as implemented in Scipy (version 0.19.1) stats module in the function ttest ind. For statistical power we used 5 seedlings, with 86 cells, including 86, 65, and 32 periclinal, longitudinal, and transverse walls respectively. Graphs were created with Python’s matplotlib (version 2.0.2, https://matplotlib.org/) pyplot module in combination with seaborn (version 0.8.0) and LaTeXs pgfplots (version 1.9, https://seaborn.pydata.org/).

## 3. Results and discussion

### 3.1. Mapping nanomechanical properties of plant cells in vivo using multifrequency AFM

The most general description of a linear viscoelastic material is given by the generalized MW model, which accounts for materials whose relaxation occurs in a set of time-scales (Section S1, SI). The storage and loss moduli, E’ and E’’, can be linked to the observables of an oscillating AFM cantilever indenting a generalized MW material by the following equations (see SI for derivation): 

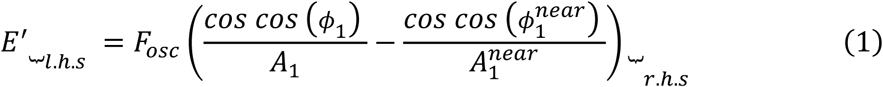

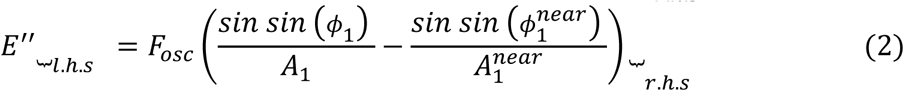

where *A*_1_ and *ϕ*_1_ are the amplitude and phase, respectively, of the cantilever. The scaling factor, 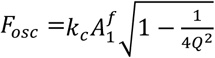 depends on the free amplitude 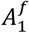 of the cantilever oscillation, the stiffness *k*_*c*_, and quality factor *Q* of the cantilever. Amplitude 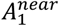 and phase 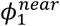, are measured at about ∼ 10 nm above the surface [4, 32] (Sections S2 and S3, SI). E’ quantifies the energy stored elastically and is related to the displacement of the material, while E’’ measures the dissipated energy and is proportional to deformation velocity. Therefore the left-hand side (l.h.s.) in Eqs. 1 and 2 depends on the viscoelastic model applied, whereas the right-hand side (r.h.s) is universal for all models. Using multifrequency AFM where the first mode of the cantilever is photo-thermally excited while feedback for imaging is done on the deflection (the zero-mode of the lever) [5] we mapped three cell junctions at the surface of the hypocotyl to simultaneously capture transverse and longitudinal anticlinal (radial), and periclinal (azimuthal) CWs (Fig. 1A,B). Using A_1_ and φ_1_ (not shown), we determined E’ and E’’, using Eqs. 1 and 2, and projected them on the topography (Fig. 1C, D). To average values for different walls, images were segmented (Section 2.5). Fig. 1E and F show significant global differences in E’ and E’’ between walls with different orientations. Although variations between periclinal and anticlinal walls probably arise from their different orientations with respect to the indentation directions, the difference in growth direction between, longitudinal and transverse walls indicates a potential link with growth-relevant mechanics. To exclude that the differences in mechanical properties in our measurements were due to the topology of the sample [19] rather than in differences in composition, we performed AFM experiments on casts of the plants made of a polymeric material (See S6 of the supplement); we did not observe differences in E’ and E’’ caused by the topology (Fig. S8). Importantly, values of E’ are very close to values obtained from non-AFM methods (see section 3.6) and thus our method overcomes the previous criticism of AFM not being capable of measuring relevant CW mechanical property values [12].

**Fig. 1:**
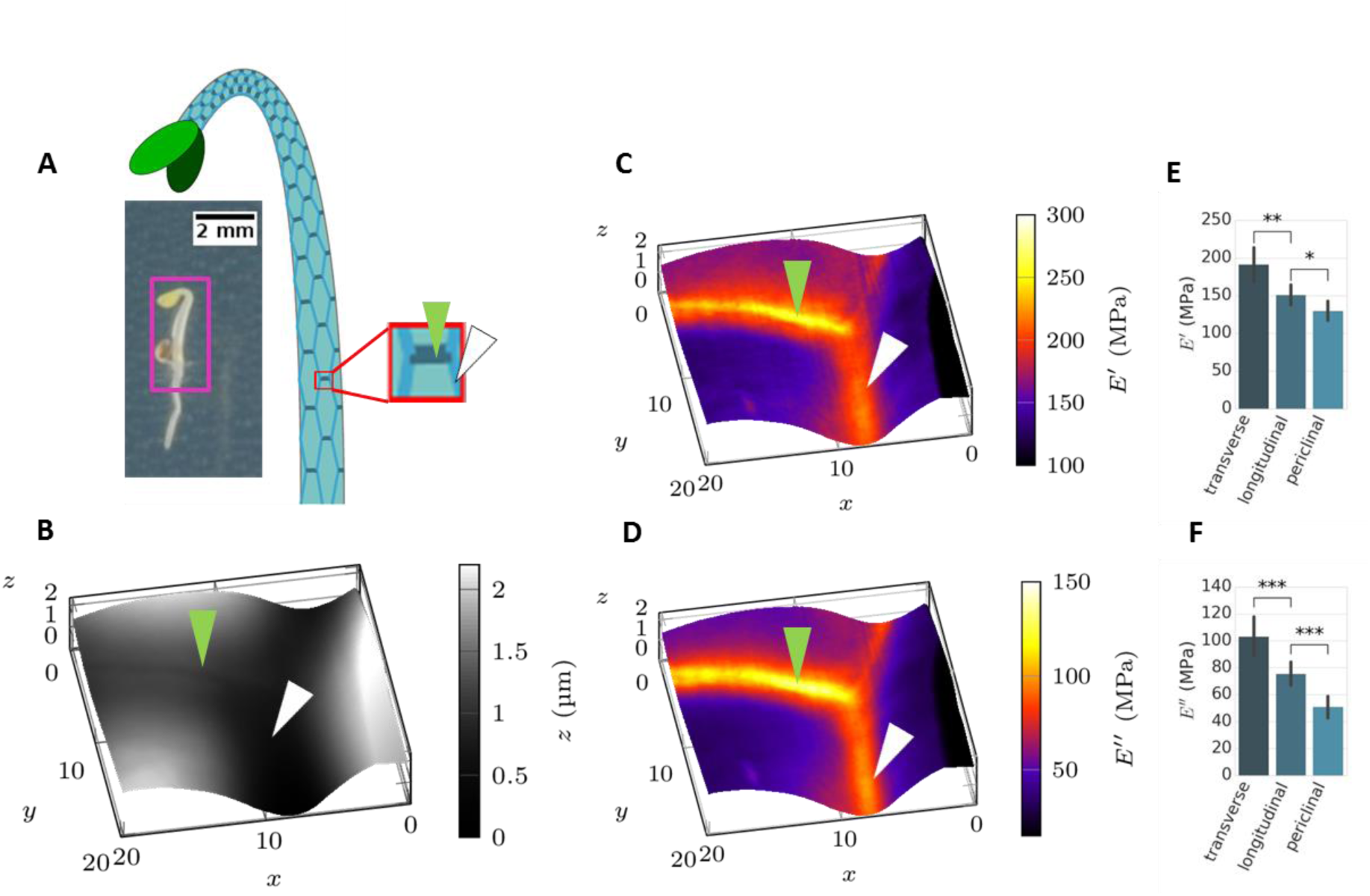
Dynamic AFM 20 µm x 20 μm images of a junction between three cells in an *Arabidopsis* hypocotyl. (A) Photograph of the seedling before imaging (scale bar: 2mm) and cartoon representing the cell structure. The colour coding for the different wall types is the same as in (E) and (F). The red frame in the cartoon represents the regions that were imaged by AFM. (B) shows an AFM topography image; the walls transverse to growth direction are marked by a green and longitudinal by a white arrowhead; the walls which are not intersections are periclinal. (C,D) maps of storage *E*^′^ and loss *E*^′′^ moduli respectively, projected onto the topography. (E, F) the statistical distributions of *E*^′^ and *E*^′′^ obtained from dynamic AFM maps of 86 cells and 4 seedlings (i.e. 86 periclinal, 65 longitudinal anticlinal, and 32 transverse anticlinal walls). * means *p* < 0.05; **, *p* < 0.01; and ***, *p* < 0.001; (Welch’s t-test). Bars are standard error of the mean.

### 3.2 Plant cell walls behave as almost perfect standard linear solid materials

To evaluate whether a given viscoelasic model can adequately explain the experimental AFM data, we developed a test that makes use of the relationship between the relaxation time-scale (τ) and the CW’s material. In the SI we show that the experimental data of E^’^ and E’’ plotted against the inverse loss tangent E’/E’’ should fit a simple mathematical function characteristic of the viscoelastic model utilized to interpret them (SI, S5). The quality of the fit can be used to determine if a model describes the material with plausible accuracy. The only assumptions made are that the material properties are continuous in adjacent pixels (it is a continuous material) and have a fixed relation between E’ and E’’. The expected behavior for the KV, MW, and SLS models if the stiffness k or the viscosity η varies is shown in Fig 2A (and Fig. S7). Fig. 2B displays a scatter plot using E’ and E’’ values obtained from Fig. 1(C,D) against E’/E’’. The graph shows an almost perfect fit of the data for all CWs to the SLS model, indicating that this model describes well the mechanical behavior of the plant cell wall. Importantly, all CWs (displayed in different colors) fit the SLS model independently of their orientation with respect to the indentation direction.

**Fig. 2:**
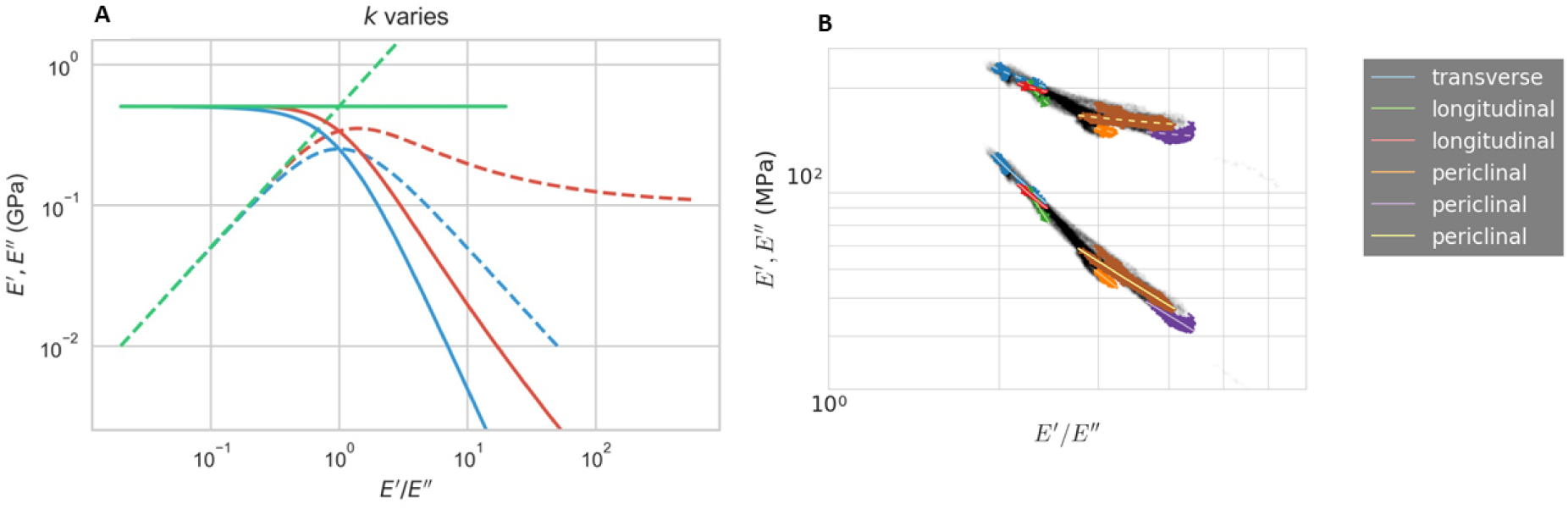
Local distribution of material properties. (A) Plot of *E*^′^and *E*^′′^ vs. the inverse loss *E*^′^/*E*′′tangent for varying *k* in the KV, MW, and SLS model (green, blue, red respectively; dashed, *E*’; solid *E*^′′^). In the SLS model *k*_*M*_ varies. (B) Scatter plot of the distribution of *E*′ and *E*^′′^ vs *E*^′^/*E*′′ for the experimental values from Fig. 1(C, D). The lines in (B) show the fit to the SLS model for constant η and changing *k*_*M*_ (dashed, *E*^′^; solid, *E*^′′^).

### 3.3 Quantitative nm resolution maps of viscosity, elasticity and relaxations times

From the fit to the SLS model (Section 3.2) values of k_∞_, k_M_, τ, η can be obtained as shown in Fig. 2C-F; k_∞_ is the relaxation spring constant, k_M_ is the MW spring constant, which relates to the initial spring constant by k_0_ = k_∞_ + k_M_, and η the viscosity, with the relation τ = η/k. Remarkably, τ values do not display significant differences depending on CW orientation (Fig. 2E), while the local values for k_∞_, k_M_, and η differ significantly for CWs of different orientations (Fig. 2D, F). Using the excitation frequency, we obtain the dimensionless quantity ωτ ≈ 1.3 which is in good agreement with theory of polymer physics [28], which predicts ωτ ≈ 1.0.

Fig. 3A – C show spatial maps of the initial spring constant k_0_ – k_M_ (1 − w_1_(κ, ωτ)), relaxation time (κ + 1)τ, and viscosity ηw_2_(κ, ωτ) (see SI,) projected on the topography of a three-cell junction. The spatial map of τ in Fig. 3 indicates that cell junctions, where more stress accumulates under turgor [39][38], have the ability to relax quicker. The mean initial stiffness k_0_ − k_M_(1 − w_1_(κ, ωτ)) is in agreement with the relaxed axial modulus previously reported[25][21]. Finally, η maps in Fig 3C indicate that viscosity varies globally, i.e. when different CWs are compared, but is relatively homogeneous locally, at the nm scale. This is confirmed in Fig. 2B where individual walls can be fitted with a single η but different η values are required for different wall types. On the other hand, the elasticity k_0_ − k_M_ (1 − w_1_ (κ, ωτ)) varies both locally and globally. Transverse walls only grow radially/circumferentially, and longitudinal walls grow both radially and longitudinally. However, hypocotyls grow substantially more in longitudinal than in circumferential or radial directions, which matches the observed pattern of viscoelastic differences between longitudinal and transverse CWs; this suggests that our technique may be suitable to measure viscoelastic properties relevant to growth.

**Fig. 3:**
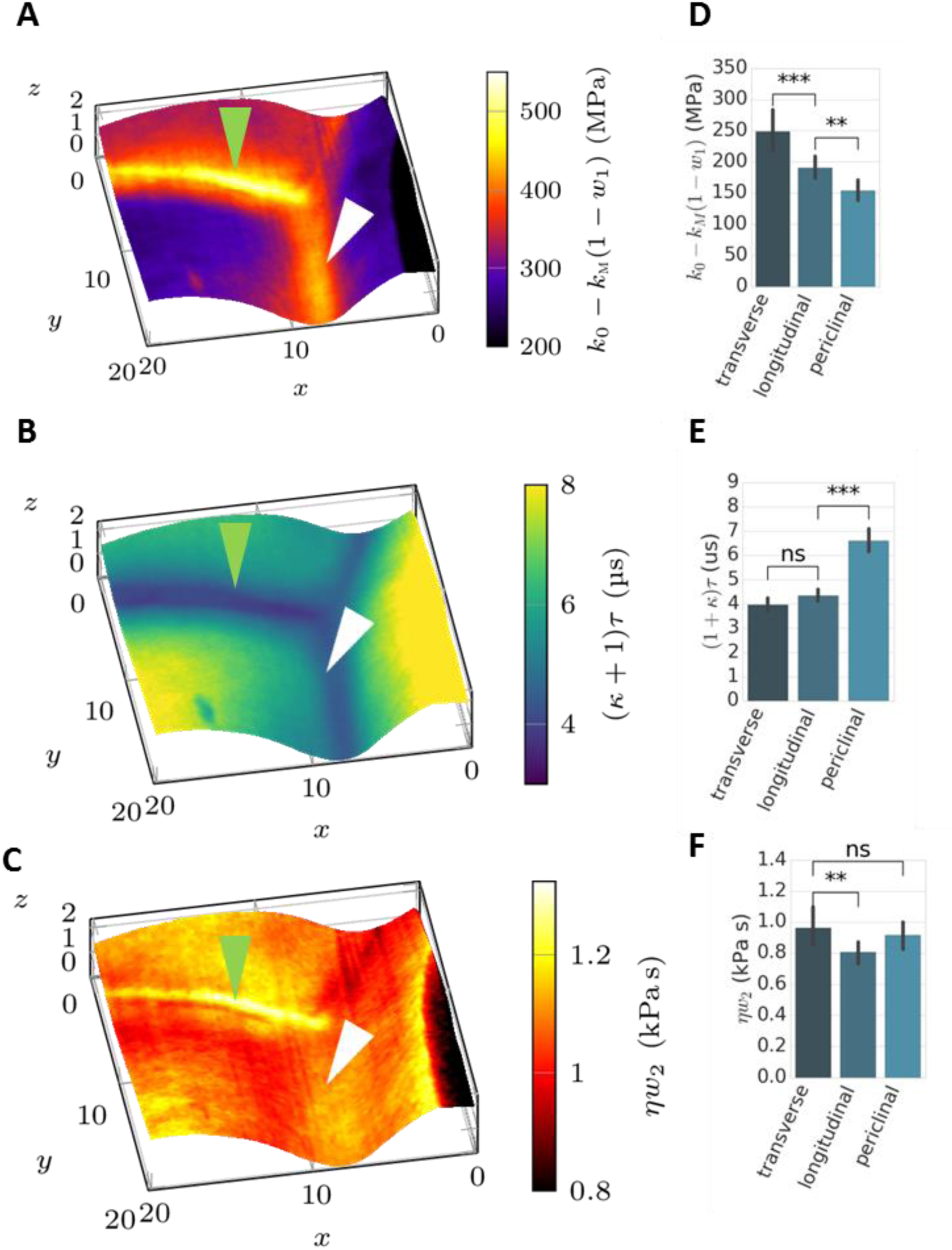
Maps of (A) the initial stiffness *k*_0_, (B) time-scale (κ + 1)τ, and (C) viscosity (κ + 1)^2^*η* calculated in the quick indentation limit and overlaid on the topography of the three cell junction shown in Fig.1A. Transverse walls are marked by green and longitudinal walls by white arrowheads. (D-F) show the statistical distribution of *k*_0_, (κ + 1)τ, and (κ + 1)^2^*η* for 86 cells from 4 seedlings; bars are standard error of the mean. Statistical significances in (D)-(F) were obtained by Welch’s t-test where ns means not significant, ** means that the p-value is < 0.01 and *** the p-value is < 0.001.

### 3.4 Nanoscale viscoelastic properties correlate to macroscopic growth speeds

As introduced in section 1, the theory of polymer dynamics confirmed by measurements of polymeric materials predicts that time scales in the millisecond range should be relevant for determining the time response of the CW at the nanometer scale [29]. Hence, we propose to test if nanoscale viscoelasticity in the millisecond time scale is related to growth speed. For testing this experimentally, we use the fact that the hypocotyl displays a growth gradient along its main axis [40-42]. We divided the hypocotyl into four regions that grow at different speeds. Fig. 4A shows the growth rate for the four zones which decreases as you go up the hypocotyl, as previously reported [40] [39]. In section S7 (SI) we show experimental data that confirm that doing AFM experiments at 60h of growth from germination we capture actively growing cells. Then, we plotted average values of E’ and E’’ against these regions. Fig. 4 shows that the lower the growth rate, the higher the values of E’ and E’’ for all CWs. Our data show that local viscoelasticity is correlated with macroscopic growth, since both E’ and E’’ are progressively higher as growth rates decrease along the hypocotyl. However, the loss tangent (related to τ as shown above) remains almost constant across the growth zones in all walls (not shown). In Fig. 4 the growth rate decreases a small amount along the hypocotyl (around 10%) while the values of E’ and E’’ increase around 70% along the same gradient. This behavior can be explained by the highly non-linear behavior of the yield threshold [6].

**Fig. 4.**
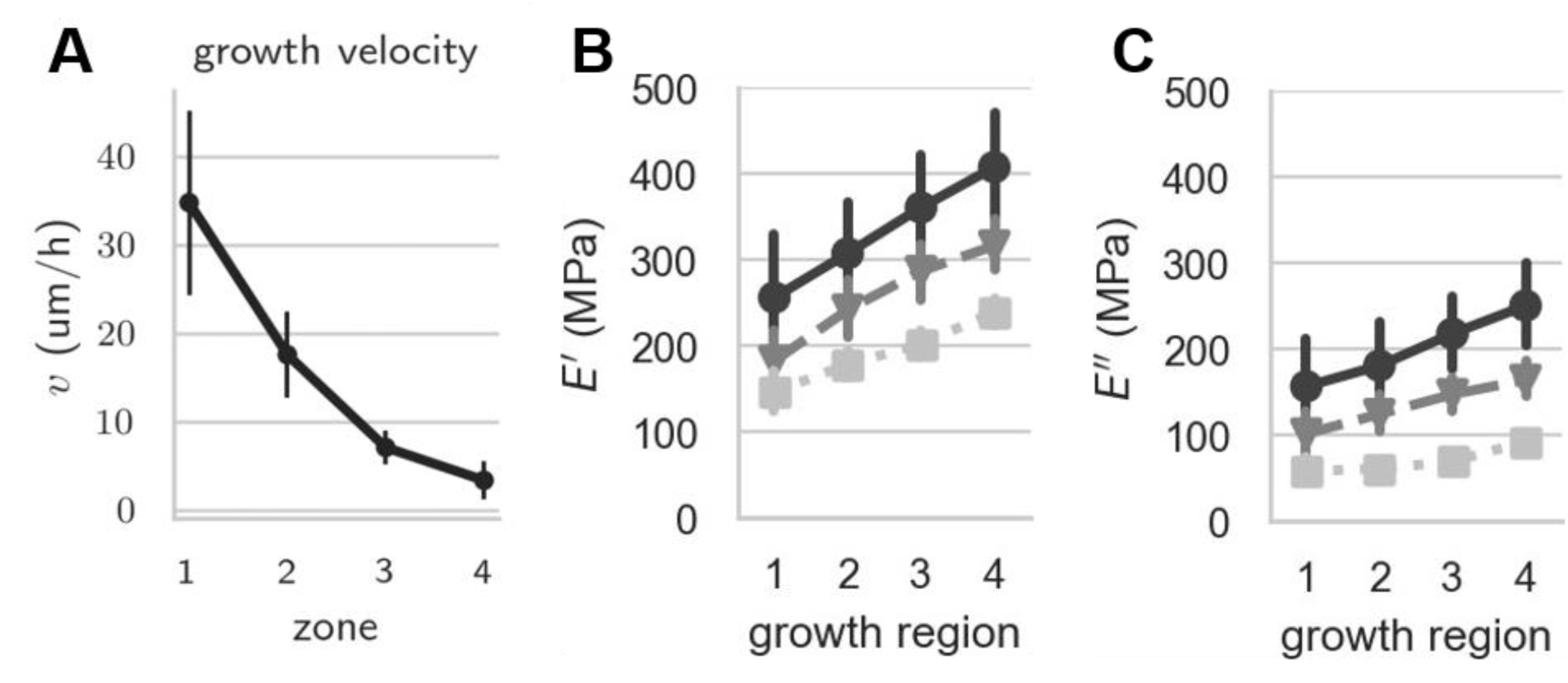
Growth gradient along the hypocotyl. (A) shows the growth rate *v* normalized against the initial cell length *L*_0_ for the 4 growth regions in the time interval 58h - 62h after transfer to 20 C from confocal microscopy image analysis (see Materials and Methods and SI). (B) and (C) show *E*^J^ and *E*^JJ^ in the same growth regions at 60h after transfer, respectively.

To investigate whether our method can measure physiologically relevant wall properties, we asked how the cellulose content of the CW changes E’ and E’’. To achieve this we added 3 nM isoxaben (IXB) to the growth medium on which the seedlings germinated and grew (see S8 of SI for details). IXB inhibits cellulose synthesis through removal of cellulose synthase complexes from the plasma membrane and therefore the total amount of cellulose in the wall is reduced [43, 44]. As a control, we measured growth rates and mechanical properties in plants grown in medium containing DMSO (see materials section). IXB-treated hypocotyls displayed a growth gradient along the hypocotyl’s longitudinal axis similar to DMSO-treated controls, with a significantly reduced cell elongation over time (Fig. S10,11). By contrast, growth in radial direction was larger or IXB treated plants, i.e. lateral/radial asymmetry is reduced (Fig. S11). After IXB treatment, both *E*′ and *E*′′ are reduced at the basal cells and as the chemically induced phenotype becomes less pronounced towards the apex, the difference between IXB-treated and control seedlings decreases (Fig. S12). The ratio of lateral to transversal quantities of E’ and E’’ (i.e. the mechanical asymmetry) was reduced (i.e., closer to one) by IXB treatment compared to DMSO controls. These results indicate that the loss of cellular asymmetry produced by IXB treatment is accompanied by a corresponding decrease in asymmetry of mechanical properties of longitudinal versus transverse walls, thereby supporting the conclusion that mechanical properties determined by dynamic AFM contain biologically meaningful information linked to cellular growth.

### 3.5 Physics of complex, hierarchical polymeric structures and growth of plant cell walls, discussion

At first sight, it seems counter-intuitive that an oscillation frequency as high as 10s of kHz would probe mechanical properties which are relevant to the time-scales of growth. However, the hierarchical nature of the structure of the cell wall (which is a confined polymer nanocomposite network) can be invoked to understand this correlation. It has been shown that long-term relaxation times of larger scales depend on short-term relaxations at smaller scales [29, 45]. The full polymer chain is characterized by Rg ∼ 10 – 100 µm whose associated time-scales are of the order of milliseconds [46]. In other words, time relaxation processes present a memory effect [45], and hence it can be expected that nm-scale properties at high frequencies can present a correlation with macroscopic properties such as growth; indeed our results experimentally demonstrate that correlation. The resonance modes of a polymer are related to its R_g_ and are dominated by the lowest mode[46]. For a typical cell of 20 µm width, it can be expected that R_g_ ∼ 10 µm; therefore the lowest mode for a polymer made only from glucose would be f_cell_ ≈ 60 kHz, which matches the magnitude of the frequencies used in our study. Our results indicate that dynamic AFM techniques such as the method presented here, as well as those reported by e.g. Raman, Contera, Cartagena et al.[4, 5, 32], Garcia, et al.[3], Proksch, et al. [47] are suitable to investigate the nanoscale mechanics underlying biological growth in vivo at relevant time-scales (milliseconds [27]) for that spatial scale.

While the correlation measured in Fig. 4 points towards a key role of the physics of polymer dynamics in the mechanical behavior of cell walls that are growing, the biological interpretation of these results is not straightforward, and requires a more sophisticated analysis. The mechanisms that allow the growth of cells rely on complex, active, dynamically controlled processes whose underlying interplay of biochemistry and physics is not well understood [48]. These processes involve dynamic remodeling of the structure in response to external conditions, and synthesis of new cell wall material, which happen at larger time scales than polymer dynamics (seconds and even minutes). However, our technique offers a new tool that can be used to disentangle the mechanical contributions of processes happening in different time scales by e.g. combining force volume methods, and dynamic high frequency methods, with external interventions such as variations of pH (which have been shown to change the mechanics of the wall by modulating the action of expansins[49] [24]). By being able to access the millisecond timescale our technique can in principle be used to disentangle pure polymer physics effects on the mechanics of the wall from other biochemical processes that work on longer time scales. Hence it can be useful to answer standing questions in the field such as “how many ways are there to loosen cell walls, to induce stress relaxation followed by cell wall growth?” [48][25]. These mechanisms are likely to involve processes at several time scales, (which are likely be nested and interdependent) and our results indeed indicate that the millisecond scale is relevant. Determining what are those mechanisms is beyond the scope of this paper, which focuses on developing a technique and demonstrating its potential use in understanding the physics of CW growth.

### 3.6 Comparison with previously reported values of elastic modulus

Previous AFM quasi-static indentation experiments of plant CWs have been put into question because they report values of elasticity (0.2 − 20 MPa range) that are orders of magnitude smaller than those obtained with other techniques such as macroscopic stretching of tissues or osmotic pressure measurements [12]. Our technique quantitatively reproduces the expected values obtained at the macro-scale and matches those of studies combining AFM with computational modelling to overcome the problems with previous semi-static AFM [24, 26]. One of these studies used CFM quasi-static indentation with a microscopic probe [24], achieving low spatial resolution, and did not produce values for E”. They reported radial moduli E_r_ = 200 − 750 MPa and axial moduli E_a_ = 120 − 220 MPa, in tobacco suspension cells, which coincides with the order of magnitude of our results. The other study used dynamic nanoindentation combined with finite element modelling to correct for geometry effects and estimate values for the elastic wall modulus ranging between 110MPa and 540MPa for young leaves [26]. The discrepancy of previous AFM results from the expected values can be explained by the fact that in a quasi-static indentation AFM experiment the bending modulus of the wall has a major contribution to the result, rather than the mechanical properties of the wall composite material. Our technique avoids this problem by imposing a 300 nm indentation to ensure a significant deformation of the CW and a contact radius of the pixel size (r_c_ ≈ 80 nm), while the 5 nm oscillations measure the actual material properties, which are obtained by the transmission of energy into the material which can be dissipated (E’’) or stored (E’). Our results qualitatively agree with semi-static indentation experiments in that longitudinal walls have lower E_j_ values than transverse walls [17]. Compared to the mechanical properties of the CW components, our values are below the range of the Young’s modulus of cellulose microfibrils, as previously demonstrated from molecular dynamic simulations and x-ray diffraction (E_transverse_ = 5 − 50 GPa, E_axial_ = 50 − 220 GPa) [39]. This suggests that the contribution from cellulose to the elasticity is reduced by the compliance of the matrix, in agreement with the prevalent current view in the field, where cellulose has the role of resisting stress elastically and pectins contribute the viscous, compliant, component [11, 15, 17].

**TABLE 1.** Average mechanical properties obtained in this paper, for comparison with values from literature.

| Mechanical properties | Transverse wall | Longitudinal wall | Periclinal wall |
| --- | --- | --- | --- |
| **$E'$ (<MPa)** | $191 \pm 12$ | $150 \pm 7$ | $129 \pm 7$ |
| **$E''$ (<MPa)** | $103 \pm 8$ | $75 \pm 4$ | $50 \pm 4$ |
| **Fit** |  |  |  |
| **$k_M$ (MPa)** | $226 \pm 19$ | $170 \pm 12$ | $110 \pm 10$ |
| **$k_\infty$ (MPa)** | $79 \pm 7$ | $71 \pm 5$ | $77 \pm 4$ |
| **$H$ (kPa s)** | $0.52 \pm 0.06$ | $0.39 \pm 0.02$ | $0.24 \pm 0.02$ |
| **$\tau$ (μs)** | $2.36 \pm 0.17$ | $2.57 \pm 0.13$ | $2.4 \pm 0.10$ |
| **MW approximation** |  |  |  |
| **$k_0 - w_1 k_M$ (MPa)** | $248 \pm 17$ | $191 \pm 9$ | $154 \pm 9$ |
| **$\eta w_2$ (kPa s)** | $0.96 \pm 0.06$ | $0.80 \pm 0.04$ | $0.91 \pm 0.05$ |
| **$\tau$ (μs)** | $3.96 \pm 0.14$ | $4.34 \pm 0.12$ | $6.58 \pm 0.25$ |

## 4. Conclusions

Novel quantitative multifrequency and other mechanical mapping AFM techniques have come forth to the microscopy field making the AFM the first tool to be able to map the mechanical properties of biological and non-biological materials with nm resolution [3, 4, 47] at high speeds. However up to now, multifrequency AFM applicability has been limited to materials that obey the Kelvin-Voigt (KV) linear viscoelastic model and hence multifrequency AFM quantitative mapping remains so far inapplicable to most real biological and biomedical cell situations in vivo, which often involve cellular growth.

Our work overcomes existing limitations: we develop a simple test to choose the correct mathematical model and a simple way to apply the correct model to AFM images to derive nm-resolution values of elasticity, viscosity, and time relaxation. We demonstrate our technique by applying it to the cells at the surface of a living growing plant, which allows us to map for the first time the viscoelasticity of the cell walls of a multicellular organism *in vivo*, including nm-maps of relaxation times of plant cell walls. Importantly for biophysics and for realizing the potential of multifrequency AFM, and establishing it as a useful tool in biology, we show that the viscoelastic nanoscale values obtained correlate with the cellular growth speeds, at 10kHz frequencies employed by the imaging method. Our results can be explained with predictions from polymer physics theory from which relaxation times at the nanoscale (which depend on the radius of gyration of polymers and are of the order of milliseconds) can be effectively probed at 10s of kHz cantilever frequencies. Our work in fact validates the applicability of AFM to study the complex multiscale mechanics of growth of multicellular organisms, providing an effective tool to probe the nanoscale. Here we have used our new method to map the living cells at the surface of Arabidopsis hypocotyls and demonstrate that the plant cell wall behaves on the nm-scale like an almost perfect SLS material. Our results complement previous studies based on mechanical stretching of whole plant organs and individual cells[10, 20, 34, 41, 42, 50], which together indicate that Arabidopsis hypocotyl CWs yield in a linear manner irrespective of length scale (nm-, micron-, or mm-scales), or the direction in which they are probed. These results suggest that CWs obey the SLS model across the scales, and raise the possibility that a biological feedback ensuring specific multiscale viscoelastic behavior is necessary for plants to maintain wild-type growth and morphology. Our results can also be put in the context of recent work that shows that cellular Gibbs energy dissipation also correlates linearly with growth in walled organisms (yeast) until it reaches saturation [46]; these results point towards a general behavior emerging from of the physics out of equilibrium that underpins growth of walled cells/organisms, where the use of metabolic energy is mirrored by that of the mechanical energy.

Importantly for the validity of our technique, our measurements of E’ = 120 MPa − 200 MPa match expected values for the plant CW unlike previous AFM quasi-static indentation experiments. Despite local patterning of km, *η*, E’, and E’’, average values of τ were not significantly different for CWs in different orientations. Interestingly, local variations in τ appeared to be affected more by alterations in elasticity, rather than viscosity (or both) which suggest a local molecular structure (i.e. cross-links density) is controlling τ (at the frequencies used in this study). Nanoscale-spatial maps of τ at cell junctions indicate that junctions can release tissue-stress faster than the purely growing regions of the periclinal walls. Additionally, average E’ and E’’ correlate with the growth rate for all CWs along the hypocotyl, indicating that both elastic and viscous contributions are crucial for plant growth across temporal and spatial scales. The fact that high frequency oscillations can pick up such correlation indicates a memory effect during cell growth as well as a fundamental polymeric relaxation behavior during cell growth. Together, our results suggest that linear viscoelasticity of plant CWs is a crucial mediator of plant growth over multiple spatial and temporal scales.

In summary, our approach expands the applicability of multifrequency AFM to deliver the quantitative time dependent nanomechanical data that are needed to understand real biological *in vivo* processes involving growth of structures based on polymer networks, including e.g. morphogenesis or tumour/biofilm progression, and more generally to correctly obtain viscoelasic properties of polymeric materials by multifrequency AFM. Our technique is particularly suitable for application to measurement of mechanics of biological epithelia, which is particularly important because the onset of 90% of cancers start with abnormal stress relaxation/accumulation patterns in epithelial cells whose quantitative study remains out of experimental reach at the nanoscale.

## Acknowledgements

In memoriam to Ian Moore. JS, SC, IM acknowledge support of the Leverhulme research project grant RPG-2014-287; JS, CK, SC, and IM were also supported by the UK Biotechnology and Biological Sciences Research Council grant BB/P010822/1. JS did the AFM experiments and the sample preparation with contributions of CK, IM and SC. CK did the confocal microscopy and determined the cell length. JS analyzed the data and prepared the graphs with contributions of SC,SK AND IM. SC and JS wrote the manuscript with contributions of IM and CK. Discussions with Antoine Jerusalem are acknowledged.

