## Supplementary material for "Mapping cellular nanoscale viscoelasticity and relaxation times relevant to growth of living Arabidopsis thaliana plants using multifrequency AFM"

### Contents:

**Sections:**

S1. A Model for Dynamic AFM Mapping of Linear Viscoelastic Materials
S2. Hydrodynamic correction
S3. Choosing the indentation model
S4. The Maxwell approximation
S5. Material Dispersion
S6. Influence of the Topography
S7. Growth rates determine the time interval of the experiments
S8. Isoxaben Treatment

**Figures:**

Figure S1: Comparison between photo-thermal and acoustic cantilever excitation in water.
Figure S2: The graphs show the behaviour of the amplitude and phase during indentation
Figure S3: The indentation curve justifies the choice of the Sneddon model.
Figure S4: Half opening angle $\alpha$ of a Nanosensors PPP-NCAuLD tip
Figure S5: Processing flowchart including the assumptions for the experimental setup and calibration as well as the assumptions necessary for calculating $E'$ and $E''$.
Figure S6: Behavior of $w_1$ and $w_2$.
Figure S7: Comparison of the material dispersion
Figure S8: Comparison of the mechanical properties of an A. thaliana hypocotyl and a latex replica of an *A. thaliana* hypocotyl.
Figure S9: A. thaliana growth rates from 48 h to 60 h and 60 h to 72 h.
Figure S10: Comparison between WT (a) and IXB (b) 3 d dark grown seedlings
Figure S11: A. thaliana growth rates from 58 h to 62 h for DMSO and IXB treatment.
Figure S12: Asymmetry of IXB treated seedlings.

### Supplementary Material

The mechanical imaging technique used in this work relies on the method that is described by Cartagena-Rivera et al. [2]. The Cartagena method is a contact resonance imaging method where the atomic force microscopy (AFM) feedback is on the deflection and at the same time the cantilever is driven at one or two oscillatory eigenmodes. However, this method is only applicable where the material can be described by the Kelvin- Voigt (KV) model, which is not accurate for the description of complex composite materials as the theory of the stress-relaxation and the creep experiments show [3]. The most general linear viscoelastic material can be described by the generalised Maxwell (MW) model which accounts for the behaviour of materials whose relaxation occurs on sev-eral timescales (Fig. 1d). This supplementary information will provide the derivation of the new theory that links the cantilever observables $A_0$, $\varphi_1$, and $A_1$, for a sinusoidal actuation to the generalised MW model. This can be reduced to the KV, MW, and standard-linear solid (SLS), which we will discuss in detail in this document. Finally, an approach to obtain the quantities $k_i$ (stiffness), $\tau_i$ (relaxation time), $\eta_i$ (viscosity) from the general MW model is developed and presented.

To obtain a relation between the cantilever observables and the mechanical properties three steps will be followed. (i) the cantilever motion is described by a driven harmonic oscillator with damping to which an external force is applied. (ii) the external force will be determined by the material parameters, which implies that the force, exerted by an oscillating generalised MW material, will be determined. (iii) the cantilever motion will be linked to an appropriate indentation model.

##### S1. A Model for Dynamic AFM Mapping of Linear Viscoelastic Materials

The Cartagena method uses the approximation of a driven harmonic oscillator moving in an external force field $F_{\text{ext}}(t)$, obtaining the following equation

$$\frac{1}{\omega_0^2}\ddot{q} + \frac{1}{\omega_0 Q}\dot{q} + q = \frac{F_{\text{osc}}\sin(\omega t)}{k_c} + \frac{F_{\text{ext}}(t)}{k_c}. \tag{S1}$$

The cantilever motion is characterised by its trajectory $q(t)$ resonance frequency $\omega_0$, spring constant $k_c$ and quality factor $Q$. The driving force oscillates with a force $F_{\text{osc}}$ and a frequency $\omega$. To solve this problem for a generalised MW material $F_{\text{ext}}(t)$ must be the resulting force of such a material. In the following derivation the external force will be denoted by $F$ instead of $F_{ext}$.

The steady state solution for $q(t)$ in eq. (S1) is given by the following expression

$$q(t) = A_0 + A_1 \sin(\omega t - \phi_1). \tag{S2}$$

Every arm in the generalised MW material is characterised by its spring constant $k_i$ and its timescale $\tau_i = \eta_i/k_i$, with the viscosity $\eta_i$. Furthermore, in the generalised MW model the total strain $\delta$ must be the same as the strain $\delta_i, i = 1 \dots N$ in every arm, i.e. $\delta_i = \delta_j \equiv \delta, \forall i, j$. The total force $F$ is given by the sum of the forces $f_i$ in every arm and a relaxation, steady state, force $f_\infty$ by

$$F = f_\infty + \sum_{i=1}^{N} f_i. \tag{S3}$$

The force $f_i$ for the MW model is given by the partial differential equation (PDE) (3)

$$k_i \frac{\partial \delta_i}{\partial t} = k_i \left( \frac{\partial \delta_{\text{dashpot}}}{\partial t} + \frac{\partial \delta_{\text{spring}}}{\partial t} \right) = \frac{1}{\tau_i} f_i + \frac{\partial f_i}{\partial t}. \tag{S4}$$

In the real space the PDEs cannot be simply summed but such an expression can be found in the Laplace domain, in which eq. (S4) has the form

$$\hat{f}_i(s) = \frac{s k_i}{s + \frac{1}{\tau_i}} \hat{\delta}_i(s). \tag{S5}$$

The Laplace transform of a function $g(t)$ is denoted by $\hat{g}(s) = \mathcal{L}(g(t))(s)$, where $s$ is the free parameter in the Laplace domain. Because of the linearity of the Laplace transform, the total $F$ can be written in the Laplace domain as

$$\hat{F}(s) = \hat{f}_\infty + \sum_{i=1}^{N} \hat{f}_i(s) = \left( \frac{\hat{f}_\infty}{s} + \sum_{i=1}^{N} \frac{sk_i}{s + \frac{1}{\tau_i}} \right) \hat{\delta}(s). \quad \text{(S6)}$$

The last equality is valid because $\delta_i = \delta, \forall i = 1 \ldots N$. With the approximation of the cantilever motion as a driven harmonic oscillator, the material stimulation can be expected to be sinusoidal. This is the case in the steady state oscillation of the driven harmonic oscillator (eq. (S2)). From this assumptions the deformation $\delta(t)$ is of the form

$$\delta(t) = \delta_0 + A_1 \sin(\omega t - \phi_1). \quad \text{(S7)}$$

In fact, in an AFM experiment the relation between the cantilever motion $\delta(t)$ and $q(t)$ is given by

$$\delta(t) = -\left(Z + q(t)\right) \quad \text{(S8a)}$$

$$\delta_0 = -\left(Z + A_0\right) \quad \text{(S8b)}$$

with the position of the $z$-piezo $Z$, and the deflection of the cantilever $A_0$.

The Laplace transform of eq. (S7) is

$$\hat{\delta}(s) = \frac{\delta_0}{s} + \frac{\omega}{s^2+\omega^2} A_1 \cos(\phi_1) - \frac{s}{s^2+\omega^2} A_1 \sin(\phi_1). \tag{S9}$$

Substituting eq. (S9) into eq. (S5) and applying the inverse Laplace transform we obtain

$$\begin{aligned} f_i(t) = &\left(k_i\delta_0 - A_1 k_i \frac{1}{1+\omega^2\tau_i^2}\left(\omega\tau_i\cos(\phi_1)+\sin(\phi_1)\right)\right) e^{-\frac{t}{\tau_i}} \\ &+ A_1 k_i \frac{\omega^2\tau_i^2}{1+\omega^2\tau_i^2}\sin(\omega t-\phi_1) \\ &+ A_1 k_i \frac{\omega\tau_i}{1+\omega^2\tau_i^2}\cos(\omega t-\phi_1). \end{aligned} \tag{S10}$$

From eq. (S10) an initial modulus $\epsilon_{0,i}$, the storage modulus $\epsilon_i'$, and the loss modulus $\epsilon_i''$ can be found as in the literature for the MW model (3)

$$\epsilon_{i,0} = k_i - \frac{1}{\delta_0\omega\tau_i}\left(\epsilon_i'\cos(\phi_1)+\epsilon_i''\sin(\phi_1)\right) \tag{S11a}$$

$$\epsilon_i' = A_1 k_i \frac{\omega^2\tau_i^2}{1+\omega^2\tau_i^2} \tag{S11b}$$

$$\epsilon_i'' = A_1 k_i \frac{\omega\tau_i}{1+\omega^2\tau_i^2}. \tag{S11c}$$

With eq. (S11) we can rewrite eq. (S10) as

$$f_i(t) = \epsilon_{0,i}\delta_0 e^{-\frac{t}{\tau_i}} + \epsilon_i' A_1 \sin(\omega t - \phi_1) + \epsilon_i'' A_1 \cos(\omega t - \phi_1)$$

$$\overset{\text{eq. (S7)}}{=} \epsilon_{0,i}\delta_0 e^{-\frac{t}{\tau_i}} + \epsilon_i'(\delta(t) - \delta_0) + \frac{\epsilon_i''}{\omega}\dot{\delta}(t). \tag{S12}$$

Using eqs. (S6) and (S10) and the linearity of the Laplace transform the total force for the generalised MW model with a relaxation spring constant $k_\infty$ is

$$F(t) = k_\infty \delta_0 + k_\infty(\delta(t) - \delta_0) + \sum_{i=1}^{N} \left( \epsilon_{0,i}\delta_0 e^{-\frac{t}{\tau_i}} + \epsilon_i'(\delta(t) - \delta_0) + \frac{\epsilon_i''}{\omega}\dot{\delta}(t) \right) \tag{S13a}$$

$$= \left( k_\infty + \sum_{i=1}^{N} \epsilon_{0,i} e^{-\frac{t}{\tau_i}} \right) \delta_0 + \left( k_\infty + \sum_{i=1}^{N} \epsilon_i' \right) (\delta(t) - \delta_0) + \left( \sum_{i=1}^{N} \epsilon_i'' \right) \frac{\dot{\delta}(t)}{\omega} \tag{S13b}$$

$$= E_0(t)\delta_0 + E'(\delta(t) - \delta_0) + E'' \frac{\dot{\delta}(t)}{\omega}. \tag{S13c}$$

In eq. (S13c) the total relaxation modulus $E_0$, storage modulus $E'$ and loss modulus $E''$ are given by the following equations:

$$E_0(t) = k_\infty + \sum_{i=1}^{N} \epsilon_{0,i} e^{-\frac{t}{\tau_i}} \tag{S14a}$$

$$E' = k_\infty + \sum_{i=1}^{N} \epsilon'_i \tag{S14b}$$

$$E'' = \sum_{i=1}^{N} \epsilon''_i . \tag{S14c}$$

Equation (S14) is well known and implemented in most finite element modelling (FEM) softwares, such as ABAQUS (11). With the solution in eq. (S13) as external force in eq. (S1) we obtain

$$\frac{1}{\omega_0^2}\ddot{q} + \frac{1}{\omega_0 Q}\dot{q} + q = \frac{F_{\text{osc}} \sin(\omega t)}{k_c} + \frac{E_0(t)\delta_0 + E'(\delta(t) - \delta_0) + E'' \frac{\dot{\delta}(t)}{\omega}}{k_c}. \tag{S15}$$

The interaction will only happen wenn the cantilever is close to the surface, and therefore the quantities in eq. (S1) must be near the surface, which will be denoted by the subscript 'near'. Substituting the steady state solution from eq. (S2) into eq. (S15), using the relations in eq. (S8) and adding in the driving force to the frequency the phase $\phi_1 - \phi_1$, i.e. $F_{\text{drive}} = F_{\text{osc}} \sin(\omega t + \phi_1 - \phi_1)$, we obtain

$$-\frac{\omega^2}{\omega^2_{\text{near}}}A_1\sin(\omega t-\phi_1)+\frac{\omega}{Q_{\text{near}}\omega_{\text{near}}}A_1\cos(\omega t-\phi_1)+A_1\sin(\omega t-\phi_1)+A_0 \tag{S16a}$$

$$=\frac{F_{\text{osc}}}{k_c}\left(\sin(\omega t-\phi_1)\cos(\phi_1)+\cos(\omega t-\phi_1)\sin(\phi_1)\right)$$
$$+\frac{1}{k_c}\left(E_0(t)\delta_0-E'A_1\sin(\omega t-\phi_1)-E''A_1\cos(\omega t-\phi_1)\right). \tag{S16b}$$

Comparing now terms with $\sin(\omega t-\phi_1)$ and $\cos(\omega t-\phi_1)$ we get the relations between the observables $k_c$, $A_1$, $\phi_1$ and the material properties

$$k_cA_0=E_0(t)\delta_0 \tag{S17}$$
$$E'=F_{\text{osc}}\frac{\cos(\phi_1)}{A_1}-k_c\left(1-\frac{\omega^2}{\omega_0^2}\right) \tag{S18}$$
$$E''=F_{\text{osc}}\frac{\sin(\phi_1)}{A_1}-k_c\frac{\omega}{Q\omega_0}. \tag{S19}$$

From Cartagena-Rivera et al. (2) we obtain the relations

$$F_{\text{osc}}=\frac{k_cA_{\text{far}}}{Q_{\text{far}}}\sqrt{1-\frac{1}{4Q^2}} \tag{S20}$$
$$1-\left(\frac{\omega}{\omega_{\text{near}}}\right)^2=\frac{F_{\text{osc}}}{k_c}\frac{\cos(\phi_{1,\text{near}})}{A_{1,\text{near}}} \tag{S21}$$
$$\frac{\omega}{\omega_{\text{near}}Q_{\text{near}}}=\frac{F_{\text{osc}}}{k_c}\frac{\sin(\phi_{1,\text{near}})}{A_{1,\text{near}}}, \tag{S22}$$

which we can substitute into eq. (S16) and the final result is

$$F_0 = k_c A_0 = E_0(t)\delta_0 \tag{S23a}$$

$$E' = F_{\text{osc}} \left( \frac{\cos(\phi_1)}{A_1} - \frac{\cos(\phi_{1,\text{near}})}{A_{1,\text{near}}} \right) \tag{S23b}$$

$$E'' = F_{\text{osc}} \left( \frac{\sin(\phi_1)}{A_1} - \frac{\sin(\phi_{1,\text{near}})}{A_{1,\text{near}}} \right). \tag{S23c}$$

In our experiments, we measured the quantities $A_{1,near}$ and $\phi_{1,near}$ near the surface, at about $\sim$ 10 nm above the sample surface. This is needed to correct for hydrodynamic effects of the liquid (1, see section S2 for details). $F_0$ is the force which is applied through the cantilever indentation and is measured by the cantilever spring constant and deflection $k_c A_0$. Equations (S23b) and (S23c) show the same relations between the cantilever observables and mechanical quantities as obtained by Cartagena-Rivera et al. (2) but the quantities $E_0(t)$, $E'$, and $E''$ are different for the different models. The reason that the relations between the mechanical properties and the cantilever observables do not change, is that $E'$ quantifies the stored energy in the system and is related to the displacement, while $E''$ is a measurement of the dissipated energy and is proportional to the velocity. Therefore, the left-hand-side (l.h.s.) in eq. (S23) will depend on the viscoelastic model applied to describe the material, whereas the right-hand-side (r.h.s.) is universal for all viscoelastic models.

For the SLS model $N = 1$, the three moduli are given as

$$E' = k_\infty + k_1 \frac{\omega^2\tau_1^2}{1+\omega^2\tau_1^2} \tag{S24}$$

$$E'' = k_1 \frac{\omega\tau_1}{1+\omega^2\tau_1^2} \tag{S25}$$

$$E_0(t) = k_\infty + \left(k_1 - \frac{A_1}{\delta_0\omega\tau_1}\left((E'-k_\infty)\cos(\phi_1) + E''\sin(\phi_1)\right)\right)e^{-\frac{t}{\tau_1}}. \tag{S26}$$

This is in agreement with the result from the literature (3).

The final contribution of the individual terms in eq. (S26) depends, eventually, on the scan speed $v$ (although $v$ is not directly involved in eq. (S26), it determines the time $t$ the cantilever spends at a pixel), the initial indentation $\delta_0$, and the oscillation amplitude $A_1$. Usually the term with $A_1/\delta_0\omega\tau_1$ is negligible, because the assumption for the derivation was $A_1 \ll \delta_0$ and for a polymer we expect that $\omega\tau_1 \sim \mathcal{O}(1)$. Furthermore, $v$ is usually that small that the time spend at a pixel $t_p \gg \tau_1$ and therefore the material is in the relaxed state and we obtain

$$E_0(t) \approx k_\infty. \tag{S27}$$

#### S2. Hydrodynamic correction

As demonstrated above it is important for quantitative dynamic measurements in liquids to correct for the hydrodynamic coupling between cantilever and surface. To keep this coupling as weak as possible we used photothermal excitation of the cantilever. Unlike acoustic excitation, which drives the whole chip of the cantilever, photothermal exci-

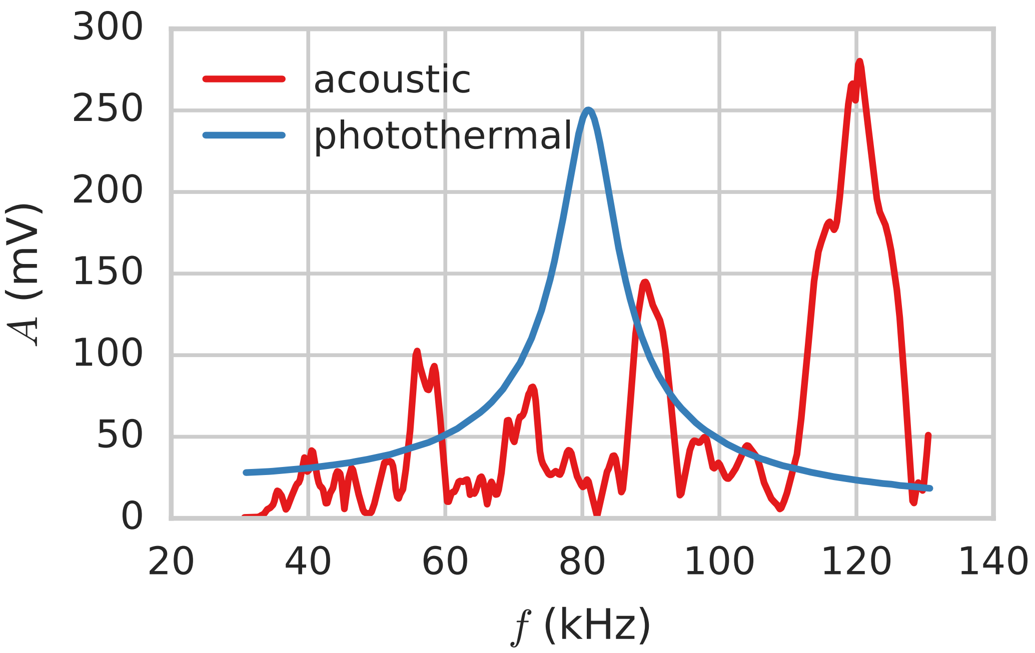


Figure S1: Comparison between photothermal and accoustic cantilever excitation in water. The advandage of photothermal excitation in water is that the resonance obeys still the laws of the harmonic oscillator and depends only weakly on the distance of the cantilever to the surface (distance dependence not shown). The multiple peaks in the case of acoustic excitation originate from the force of a water wave interacting with the cantilever. The water wave is caused by the oscillation of the cantilever chip.

tation drives only the cantilever and consequently the excitation of the water is reduced by the ratio of the surface area of the two. This makes the oscillation in water small and therefore, the cantilever response is linear (cf. fig. S1). This simple response allows a correction for the hydrodynamic coupling. It should be mentioned here that any direct excitation of the cantilever is possible. The Lorentz force excitation as used by Raman et al. (10) requires special cantilevers, which are only produced

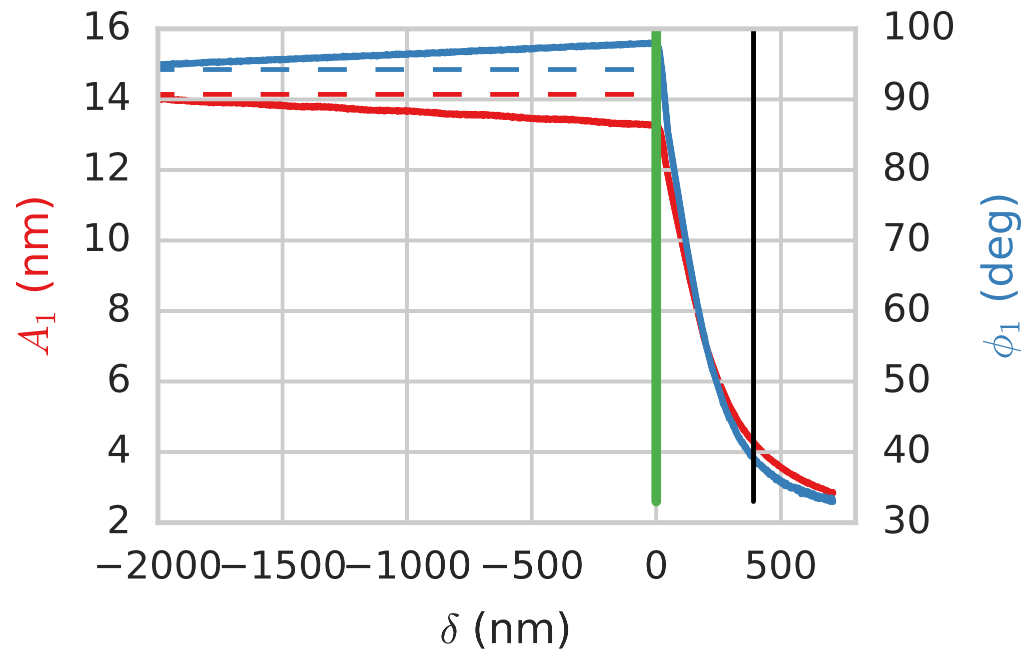


Figure S2: The graphs show the behaviour of the amplitude and phase during indentation. The green line shows the contact point of the force-distance curve (cf. fig. S3) and the dashed/solid lines show the free amplitude and phase far away from the surface and close to the surface, respectively.

in limited varations. Due to the low stiffness of these cantilevers it was not possible to use them for our experiment. Therefore, due to the complex topology of multicelluar plant tissues compared to isolated cells, high aspect ratio cantilevers are required to avoid contact between the cantilever and the tissue. Sich cantilevers do not exist for Lorentz force excitation.

Having a system were a hydrodynamic correction is possible makes this an easy procedure. A force-distance curve has to be measured while the cantilever is oscillating and this gives the phase and amplitude just above the surface (cf. fig. S2). Usually, we chose a distance about 10 nm above the surface and for the free amplitude and phase the first value of those curves was chosen. As mentioned above the tissue topography can influence the measurements and this is not only true for the physical contact between sample and cantilever but also for the hydrodynamic coupling. This means the hydrodynamic coupling will increase the more area of the cantilever is close to the surface and for more accurate results this effect had to be corrected by an additional measurement, i.e. repeating the correction measurement. The first line would measure the topography and mechanical interaction and the second measurement has to be just above the surface to see how the hydrodynamic coupling changes. However, since the coupling is weak and the correction is done for all data in the same way it will not have any consequences for the interpretation and the behaviour but only for the exact mechanical quantities. Using

$$E' = F_{\mathrm{osc}} \left( \frac{\sin(\phi)}{A} - \frac{\sin\left(\tilde{\phi}\right)}{\tilde{A}} \right)$$
$$E'' = F_{\mathrm{osc}} \left( \frac{\cos(\phi)}{A} - \frac{\cos\left(\tilde{\phi}\right)}{\tilde{A}} \right)$$

and assume $\Delta x_i = \xi x_i$ the error can be estimated to

$$\begin{aligned}
\Delta E' &= \sqrt{\sum_i \left(\frac{\partial E'}{\partial x_i}\Delta x_i\right)^2} \\
&= F_{\text{osc}}\sqrt{\left(\frac{\Delta \sin(\phi)}{A}\right)^2 + \left(\frac{\sin(\phi)\Delta A}{A^2}\right)^2 + \left(\frac{\Delta \sin\left(\tilde{\phi}\right)}{\tilde{A}}\right)^2 + \left(\frac{\sin\left(\tilde{\phi}\right)\Delta \tilde{A}}{\tilde{A}^2}\right)^2} \\
&= F_{\text{osc}}\xi\sqrt{2}\sqrt{\left(\frac{\sin(\phi)}{A}\right)^2 + \left(\frac{\sin\left(\tilde{\phi}\right)}{\tilde{A}}\right)^2}.
\end{aligned}$$

Since

$$\left(\frac{\sin(\phi)}{A}\right)^2 + \left(\frac{\sin\left(\tilde{\phi}\right)}{\tilde{A}}\right)^2 = \frac{E'^2}{F_{\text{osc}}^2} + 2\frac{\sin(\phi)\sin\left(\tilde{\phi}\right)}{A\tilde{A}}$$

the relative error is

$$\frac{\Delta E'}{E'} = \sqrt{2}\xi\sqrt{1 + \frac{F_{\text{osc}}^2}{E'^2}\frac{\sin(\phi)\sin\left(\tilde{\phi}\right)}{A\tilde{A}}} \leq 2\xi.$$

The inequality is true because $F_{\text{osc}}^2 = \mathcal{O}\left(E'^2 A\tilde{A}\right)$. Assuming $\xi \approx 0.05$, which is a reasonable assumption for our experiment, then the relative error is $\Delta E'/E' \leq 10\,\%$. The same derivation works also for $E''$

$$\frac{\Delta E''}{E''} = \sqrt{2}\xi\sqrt{1 + \frac{F_{\text{osc}}^2}{E''^2}\frac{\cos(\phi)\cos\left(\tilde{\phi}\right)}{A\tilde{A}}} \leq 2\xi.$$

#### S3. Choosing the indentation model

To link the viscoelastic values from the one dimensional standard linear solid model, which was used in the derivation above, to three dimensional moduli an appropriate indentation model has to be chosen. This can be very complex since the plant cell wall is an intricate composite material. However, we can assume that at small scales, such as the size of the volumes at which the indentation takes place, the sample behaves isotropically. Additionally, the wall is much thicker than the oscillatory indentation depth, which is about $\sim 5\,\mathrm{nm}$, that we can assume that the major contribution comes from indenting the material rather than bending it. With the assumptions of local isotropy and having an infinitely thick material all the models from indentation theory can be used. For linear elasticity that implies

$$F = g\frac{E}{1-\nu^2}\delta^\gamma, \tag{S28}$$

where $\nu$ is the Poisson ratio of the sample and $E$ the corresponding Young's modulus, and $g$ and $\gamma$ are geometrical coefficients specific to the indenter geometry.

For the Sneddon model as used in the fit, which is shown in fig. S3 $g = 2\tan(\alpha)/\pi$ with $\alpha \approx 15°$ being the half opening angle, which is given by the manufacturer of the cantilever (cf. fig. S4) and $\gamma = 2$ (12).

From this model the three dimensional (3D) moduli can be determined by following the approach of Raman et al. (10) and Cartagena-Rivera et al. (2). With a Taylor series expansion of the indentation model in eq. (S28) we obtain

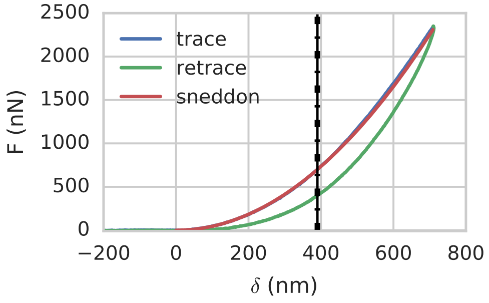


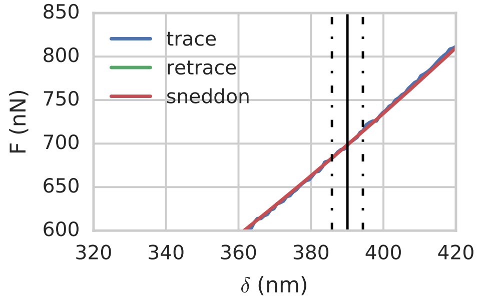


Figure S3: The indentation curve justifies the choice of the Sneddon model as the fit shows. The image on the r.h.s. shows a zoom from the l.h.s. image. The solid black line shows the indentation depth and the dashed line show the oscillation amplitude.

$$F(\delta) = F(\delta_0) + \frac{\partial F}{\partial \delta}(\delta - \delta_0) + \frac{\partial F}{\partial(\omega\delta)}\dot{\delta} \tag{S29}$$

$$= g\frac{E_0(t)}{1-\nu^2}\delta_0^\gamma + \gamma g \frac{E'_{3D}}{1-\nu^2}\delta_0^{\gamma-1}(\delta - \delta_0) + \gamma g \frac{E''_{3D}}{1-\nu^2}\delta_0^{\gamma-1}\frac{\dot{\delta}}{\omega}. \tag{S30}$$

The introduction of $E'_{3D}$ and $E''_{3D}$ is reasoned by the knowledge we deduced from the one dimensional (1D) derivation that $E'$ is related to the displacement $(\delta(t) - \delta_0)$ and $E''$ is related to the velocity $\dot{\delta}(t)$ (eq. (S13)).

Comparing eq. (S30) with eq. (S13) we find

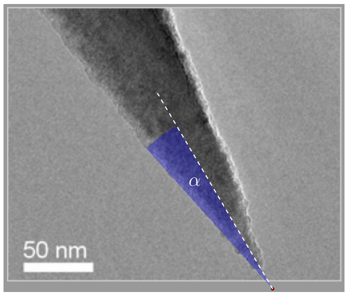


Figure S4: Half opening angle $\alpha$ of a Nanosensors PPP-NCAuLD tip (8). The angle has to be measured for a virtual tip, which is outside the real probe since the real tip size is finite. However, the fit in fig. S3 demonstrates that the experiment behaves according to this simplified model, justifying the choice of the model.

$$F(\delta_0) = F_0 = g\frac{E_0(t)}{1-\nu^2}\delta_0^\gamma \tag{S31a}$$

$$E'_{1D} = \gamma g \frac{E'_{3D}}{1-\nu^2}\delta_0^{\gamma-1} \tag{S31b}$$

$$E''_{1D} = \gamma g \frac{E''_{3D}}{1-\nu^2}\delta_0^{\gamma-1}. \tag{S31c}$$

From here the three 3D moduli can be obtained by

$$\delta_0 = \sqrt[\gamma]{\frac{F_0}{E_0}\frac{1-\nu^2}{g}} \tag{S32a}$$

$$E'_{3D} = \frac{E'_{1D}}{\gamma}\sqrt[\gamma]{\frac{1-\nu^2}{g}}\sqrt[\gamma]{\frac{E_0}{F_0}}^{\gamma-1} \tag{S32b}$$

$$E''_{3D} = \frac{E''_{1D}}{\gamma}\sqrt[\gamma]{\frac{1-\nu^2}{g}}\sqrt[\gamma]{\frac{E_0}{F_0}}^{\gamma-1}. \tag{S32c}$$

$E_0$ can be determined from the quasi-static indentation as shown in fig. S3 and with the assumption of eq. (S27) $E_0$ is the relaxation modulus, which we assume to be constant for the whole sample. We then obtain $F_0, E'_{1D}, E''_{1D}$ from the AFM scan. Furthermore we assume the cell wall (CW) is an incompressible material and therefore $\nu = 0.5$. Moreover, fig. S3 shows that our indentation follows the Sneddon model (left panel), i.e. $\gamma = 2, g = \frac{2}{\pi}\tan(\alpha)$, and that the experiment was done in the limit of small oscillations where the linear approximation fits very well (right panel). With this experimental evidence we obtain from eq. (S32)

$$E'_{3D} = \frac{E'_{1D}}{2}\sqrt{\frac{1-\nu^2}{\tan(\alpha)}\frac{\pi}{2}\frac{E_0}{F_0}} \tag{S33a}$$

$$E''_{3D} = \frac{E''_{1D}}{2}\sqrt{\frac{1-\nu^2}{\tan(\alpha)}\frac{\pi}{2}\frac{E_0}{F_0}}. \tag{S33b}$$

It is important to highlight, that Cartagena-Rivera et al. (2) chose $E'_{3D}$ instead of $E_0$ because they use the KV model. This approach overestimates the material stiffness because it neglects the relaxation which happens throughout the scan in contact mode. Comparing the mathematical expressions from eq. (S33) with the result of Cartagena-Rivera et al. (2) $E'_{3D} = \pi(1-\nu^2){E'}^2_{1D}/(8\tan(\alpha)F_0)$, demonstrates that in the overestimation of the KV is given by a factor

$$\frac{{E'}^{\mathsf{GMW}}_{3D}}{{E'}^{\mathsf{KV}_{3D}}} = \sqrt{\frac{1-\nu^2}{\tan(\alpha)}\frac{\pi}{2}}\frac{E'_{1D}}{\sqrt{F_0E_0}} \approx 1.3\frac{E'_{1D}}{\sqrt{F_0E_0}}.$$

The same relation can be found for the loss modulus. With a relax-

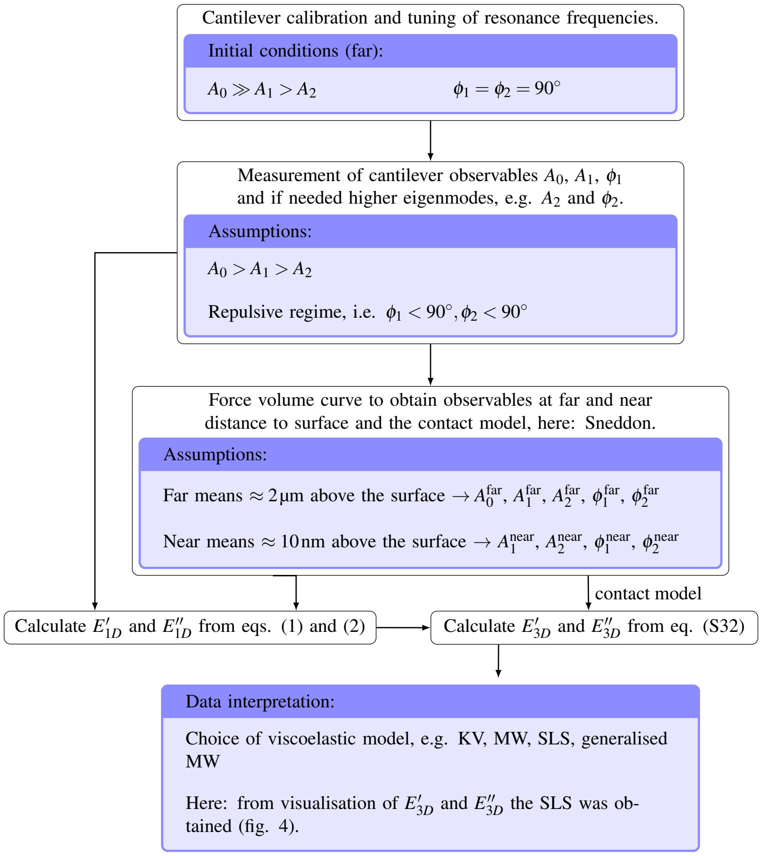


Figure S5: Processing flowchart including the assumptions for the experimental setup and calibration as well as the assumptions necessary for calculating $E'$ and $E''$.

ation modulus of $E_0 \approx 30$ MPa and an applied force of $F_0 \approx 750$ nN and $E'_{1D} \approx 40$ N m$^{-1}$ the KV model provides values which are about a factor 4.25 larger than the generalised MW model.

To summarise, in order to obtain dynamic viscoelastic maps the following conditions and experimental order must be fulfilled:

- $A_0 \gg A_1 > A_2$ to ensure linearisation of the problem, i.e. contact area does not change significantly during oscillation (see also fig. S3)

- The experiment must take place in the repulsive regime, i.e. $\phi_1 < 90°$ and $\phi_2 < 90°$
- Force-volume curves must be obtained after every scan and record the parameters $A_0$, $A_1$, $\phi_1$, and if measured in the experiment also higher eigenmodes. The calibration quantities near the surface should be measured at about 10 nm above the surface and the quantities far from the surface should be measured at least 2 µm above the surface. This curve is also important to determine the contact model, i.e. exponent $\gamma$ in eq. (S28).
- With the scan and the calibration quantities $E'_{1D}$ and $E''_{1D}$ can be calculated.
- With the contact model and the fitted Young's modulus $E_0$ from this model and the one dimensional quantities $E'_{3D}$ and $E''_{3D}$ can be determined.
- The choice of the viscoelastic model, e.g. MW, KV, SLS, is important for the data interpretation but not for the experiment.

The whole process, including mathematical and experimental conditions is summarized in a flowchart (fig. S5).

**S4. The Maxwell approximation**

From eqs. (S24) and (S25) we obtain

$$\frac{E'}{E''} = \frac{\kappa}{\omega\tau} + (1+\kappa)\omega\tau \equiv \Theta(\kappa, \omega\tau). \tag{S34}$$

We call $\Theta(\kappa, \omega\tau)$ the time response function and we introduced $\kappa = k_\infty/k_{\mathsf{M}}$. For the MW model we obtain the relations for $k_{\mathsf{M}}$ and $\eta$

$$k_{\mathsf{M}} = \frac{{E'}^2 + {E''}^2}{E'} \tag{S35a}$$

$$\eta = \frac{1}{\omega}\frac{{E'}^2 + {E''}^2}{E''}. \tag{S35b}$$

Using the approach at the r.h.s. of eq. (S35) we obtain for the SLS model

$$\begin{aligned}
\frac{{E'}^2 + {E''}^2}{E'} &= E' + \frac{E''}{\Theta(\kappa, \omega\tau)} && \text{(S36a)}\\
&= k_0 - k_{\mathsf{M}} + k_{\mathsf{M}}\frac{\omega^2\tau^2}{1+\omega^2\tau^2} + k_{\mathsf{M}}\frac{\omega\tau}{1+\omega^2\tau^2}\frac{1}{\Theta(\kappa, \omega\tau)}\\
&= k_0 - k_{\mathsf{M}}\left(1 - \frac{\omega^2\tau^2}{1+\omega^2\tau^2}\left(1 + \frac{1}{\Theta(\kappa, \omega\tau)\omega\tau}\right)\right)\\
&\equiv k_0 - k_{\mathsf{M}}\left(1 - w_1(\kappa, \omega\tau)\right)\\
\frac{1}{\omega}\frac{{E'}^2 + {E''}^2}{E'} &= E''\left(1 + \left(\frac{E''}{E'}\right)^2\right) && \text{(S36b)}\\
&= \eta\frac{1+\Theta(\kappa, \omega\tau)^2}{1+\omega^2\tau^2} \equiv \eta w(\kappa, \omega\tau), && \text{(S36c)}
\end{aligned}$$

where we introduced the weight factors $w_1(\kappa, \omega\tau), w_2(\kappa, \omega\tau)$

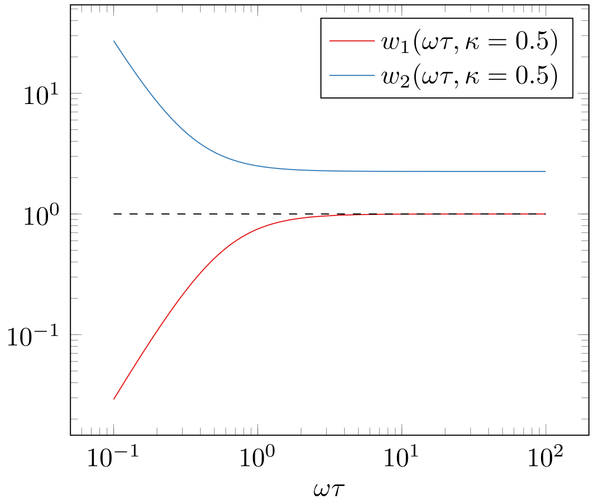


Figure S6: Behavior of $w_1$ and $w_2$.

$$w_1(\kappa, \omega\tau) = \frac{\omega^2\tau^2}{1+\omega^2\tau^2}\left(1 + \frac{1}{\Theta(\kappa,\omega\tau)\omega\tau}\right) \quad \text{(S37a)}$$

$$w_2(\kappa, \omega\tau) = \frac{1+\Theta(\kappa,\omega\tau)^2}{1+\omega^2\tau^2}. \quad \text{(S37b)}$$

Both $w_1$ and $w_2$ depend on the significant timescale $\omega\tau$ but $w_1 \in [0, 1]$ introduces a reduction of $k_0 = k_\infty + k_{\mathsf{M}}$ by a percentage of $k_{\mathsf{M}}$, whereas $w_2 \geq 1$ introduces an additional resistance which arises from the presence of $k_\infty$, which can be seen by the consideration of the equality $w_2 = 1 \Leftrightarrow \kappa = 0 \Rightarrow k_\infty = 0$.

#### S5. Material Dispersion

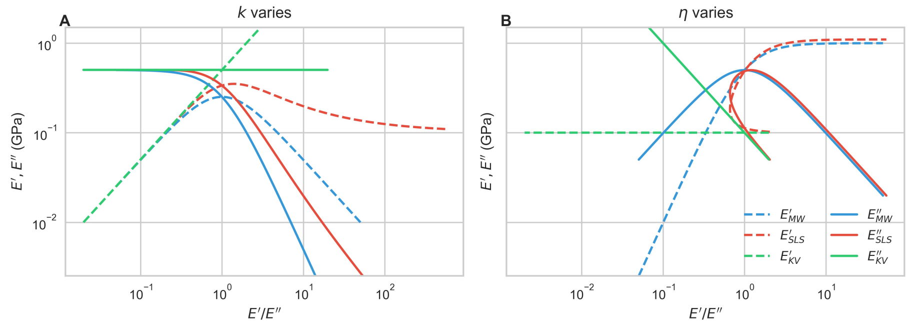


Figure S7: Comparison of the material dispersion. **a**, only $k$ changes (as in main article) and **b** only $\eta$ changes

#### S6. Influence of the Topography

A frequently discussed issue is whether the topography of the cells influences the mechanical properties. This hypothesis was tested using a latex replica of the hypocotyl. If the characteristic distribution in the mechanical mapping in fig. S8 (a) is caused by the topography, the mechanical profile of the replica (cf. fig. S8 (c)) should qualitatively have the same distribution. However, this is not the case and suggests that the hydrodynamic effect is not responsible for the mechanical behaviour of the plant samples at the cell junctions but it is caused by differences in the mechanical properties or wall internal stresses.

##### *S6.1. Preparation of Hypocotyl Casts and Moulds*

To investigate topographical contributions during mechanical mapping it was useful to create replicas of the sample as a control, made from less complex materials in order to asses the contribution of the topography alone. The existing protocols Dumais and Kwiatkowska (4)

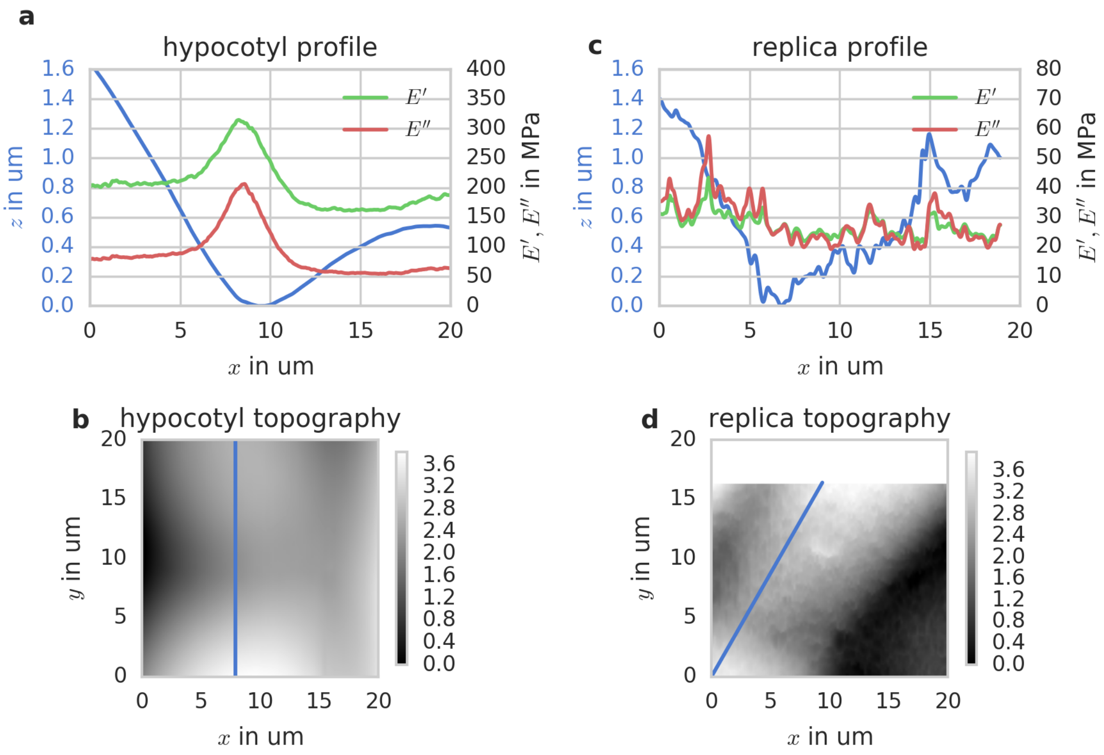


Figure S8: Comparison of the mechanical properties of an *A. thaliana* hypocotyl and a latex replica of an *A. thaliana* hypocotyl. ( (a)) shows the line profile of the topography of the seedling, which is indicated by the blue line in ( (b)), and the corresponding mechanical properties. ( (c)) shows the line profile of the topography of the replica, which is indicated by the blue line ( (d)), and the corresponding mechanical properties.

turned out to not being useful for the application of viscoelasatic mapping, because dental waxes were too stiff for the investigation of dynamic mechanical properties. Instead we used a polymorphic plastic compound from ThermoMorph® to create the mould. The plastic was purchased via amazon.co.uk.

The Thermomorph was heated in water to 70 °C. In this state this substance was soft enough to be deformed easily by the seedlings. For a high quality cast the plastic must be spread evenly on a substrate. It was important to achieve a smooth surface and at the same time the compression of the material should not create domains in deeper layers because this causes a resistance when the seedling was pressed into the material. To avoid these artefacts the plastic was formed to a small ball of a diameter $d \approx 5\,\mathrm{mm}$ using thumb and index finger. The exact size depended on the size of the seedling. In this project seedlings of an age of about 3 d after transfer to the culture room were used.

The plastic ball was then compressed between two glass slides into a thin layer and after the layer had the desired dimensions the top slide is removed and the seedling is pressed into the layer using a slide. After the material became rigid the slides and the seedling were removed. During the whole process it was useful to keep the hydration layer of the plastic. This helped when removing the slides and the seedling which are hydrophilic whereas the plastic is hydrophobic.

The cast were filled with liquid moulding latex from the Vesey gallery in Birmingham, UK. The amount required depended on the size of the seedling. However, it was important that not only the cast was filled but also that a little bit of the smooth surface was covered. This made it easier to remove the mould from the cast and attach it to a sample holder after the latex dried. If it was difficult to separate cast and mould from each other the cast could be covered in a layer of sunflower oil or any other surfactant. But the droplet formation of the surfactant could create artefacts in the mould and therefore a uniform monolayer must be achieved.

Before the mould could be attached to the sample holder, unnecessary parts were removed with a razor blade. This made it easier to find the replica on the latex surface. The mould can be attached with any adhesive to the holder. We used the medical adhesive, which was also used to attach the seedlings.

#### S7. Growth rates determine the time interval of the experiments

We chose to perform experiments on dark-grown hypocotyls, which are known to display a developmental gradient along their longitudinal axis (5). Basal cells expand first, with a wave of growth subsequently shifting in acropetal direction. To quantify cellular growth rates in different regions of the hypocotyl under our growth conditions, we examined growth rates in two time intervals: 48 h to 60 h and 60 h to 72 h after shifting plants to the growth chamber. Figure S9 shows the average hourly growth rate between 48 h to 60 h and 60 h to 72 h old seedlings after ger-

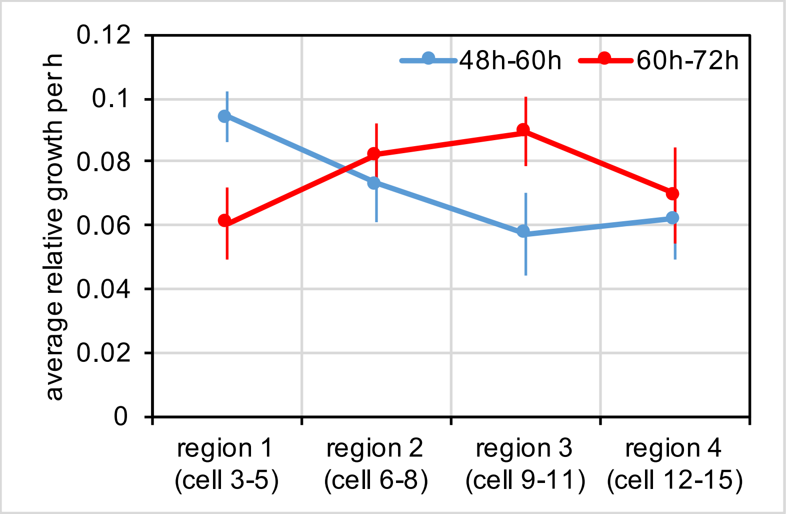


Figure S9: *A. thaliana* growth rates from 48 h to 60 h and 60 h to 72 h. The error bars represent standard deviation.

mination. It can be clearly seen the two intervals display a significantly different growth behaviour recapitulating the acropetal wave of growth described in the literature. In the first time interval, cells at the base displayed the most rapid growth, whereas this growth peak had shifted towards zone 3 in the second time interval. We performed our AFM experiments at 60h to capture actively growing cells at different regions 1 to 4. Based on the assumption that cell wall mechanical properties undergo significant changes to drive the observed shift in growth rate along the hypocotyl in the time scales observed here, we subsequently focussed on a narrower time interval (58 h to 62 h) to quantify growth rates relevant to our AFM mechanical measurements.

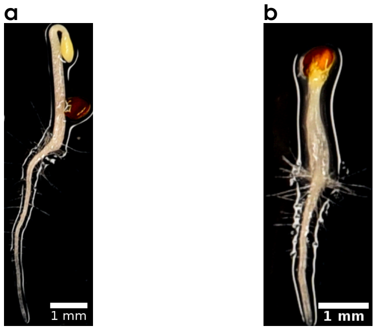


Figure S10: Comparison between WT (a) and IXB (b) 3 d dark grown seedlings.

#### S8. Isoxaben Treatment

To investigate whether our method can measure physiologically relevant wall properties, we asked how the cellulose content of the CW changes its mechanical properties. To achieve this we added 3 n isoxaben (IXB) to the growth medium on which the seedlings germinated and grew them for 3 d (see materials and methods). IXB inhibits cellulose synthesis through removal of cellulose synthase complexes from the plasma membrane (9) and therefore the total amount of cellulose in the wall is reduced (6). The hypocotyls show a phenotype of severe cell swelling and reduced organ growth (fig. S10 (b)), the latter of which is an active response to cellulose deficiency (7). Although the swelling phenotype reduces from the basal part towards the apex of the hypocotyl corresponding to the growth gradient along the hypocotyl, the IXB treated seedlings do not form an apical hook , which depends on differential cell elongation at the apex of the hypocotyl. We quantified cellular growth rates in IXB and dimethyl sulfoxide (DMSO) treated hypocotyls, and found that although IXB-treated hypocotyls still displayed a growth gradient along the hypocotyl's longitudinal axis similar to DMSO-treated controls, cell elongation was significantly reduced (fig. S10). By contrast, growth in radial direction was significantly increased. The resulting shift towards more isotropic growth at the cellular level matches the observed macroscopic phenotype. To test whether this shift in cellular growth pattern was reflected in the cells mechanical properties, we measured. With reduced cellulose content one would not only expect the cells to swell but also a reduction in their elastic modulus. Figure S12 shows $E'$ and $E''$ for 9 IXB treated and 6 DMSO treated control seedlings. The dots represent the mean values of 224 and 176 cells for treatment and control, respectively, including 77 and 58 transverse, 150 and 116 longitudinal, and 224 and 176 periclinal walls. The error bars represent the 1000 fold bootstrapped error of the mean with a 95 % confidence interval. The different zones are counted from the base (1) towards the apical hook (4) of the hypocotyl to create an average over several cells of the same seedling. Figure S12 shows clearly, that after IXB treatment, both $E'$ and $E''$ are reduced at the basal cells and as the chemically induced phenotype becomes less pronounced towards the apex the difference between treated and control seedlings decreases. To examine whether the reduced anisotropy in growth was reflected in a shift in asymmetry of mechanical properties in transverse vs longitudinal walls, we calculated asymmetry in storage and loss modulus as well as growth (fig. S13). For all measures, asymmetry was reduced (i.e., closer to one) by IXB treatment compared to DMSO controls. Therefore, we can demonstrate that the loss of growth asymmetry through IXB treatment is indeed linked to a loss in asymmetry of mechanical properties at longitudinal versus transverse walls, supporting the notion that mechanical properties determined by dynamic AFM can be related to cellular growth rate. It should be noted that absolute values for storage and loss moduli do not correlate with growth rate across both treatments – although both storage and loss moduli are substantially reduced in IXB-treated seedlings (presumably due to the reduction in stiff cellulose), this does not translate into increased growth rates. Based on previous observations that plants can detect cellulose deficiencies and actively reduce cellular growth rates (7), this observation is not surprising, and supports the notion that regulation of growth is a complex, multifactorial process. Nevertheless, our IXB experiment demonstrates that mechanical properties as measured by AFM contain biologically meaningful information linked to cellular growth.

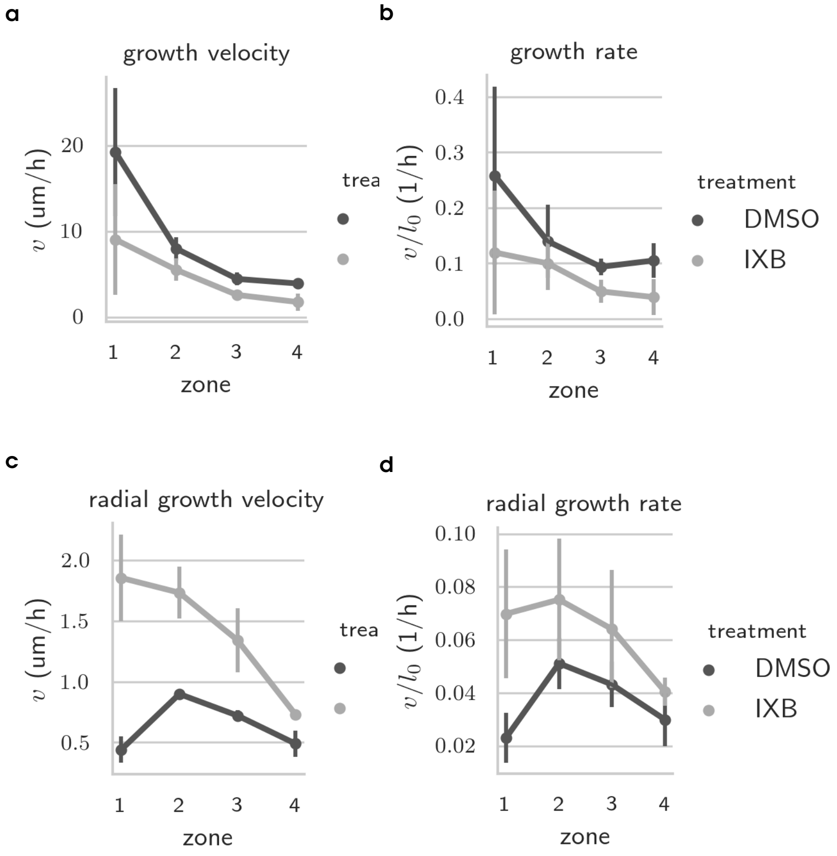


Figure S11: *A. thaliana* growth rates from 58 h to 62 h for DMSO and IXB treatment.

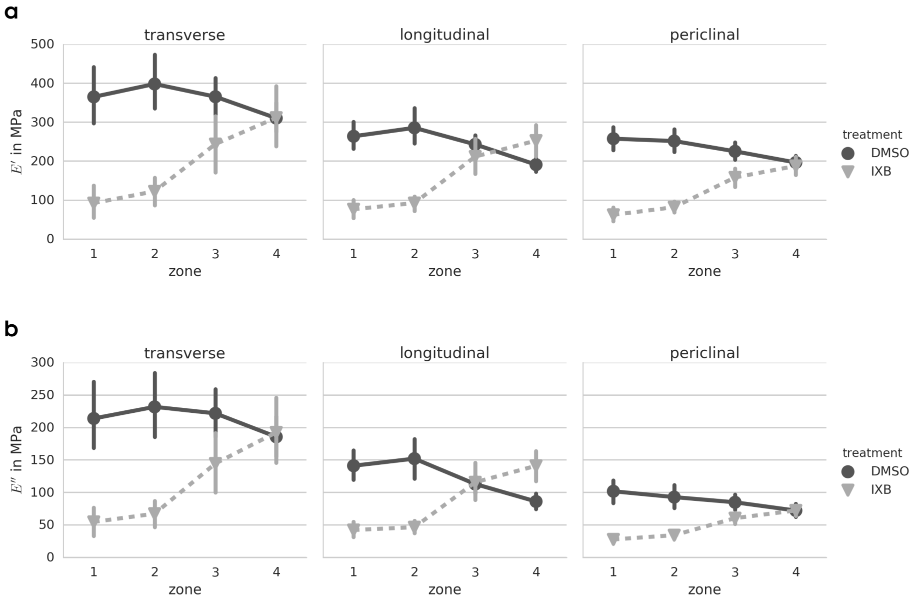


Figure S12: Viscoelasticity of IXB treated seedlings. Maps of storage (a) and loss (b) moduli.

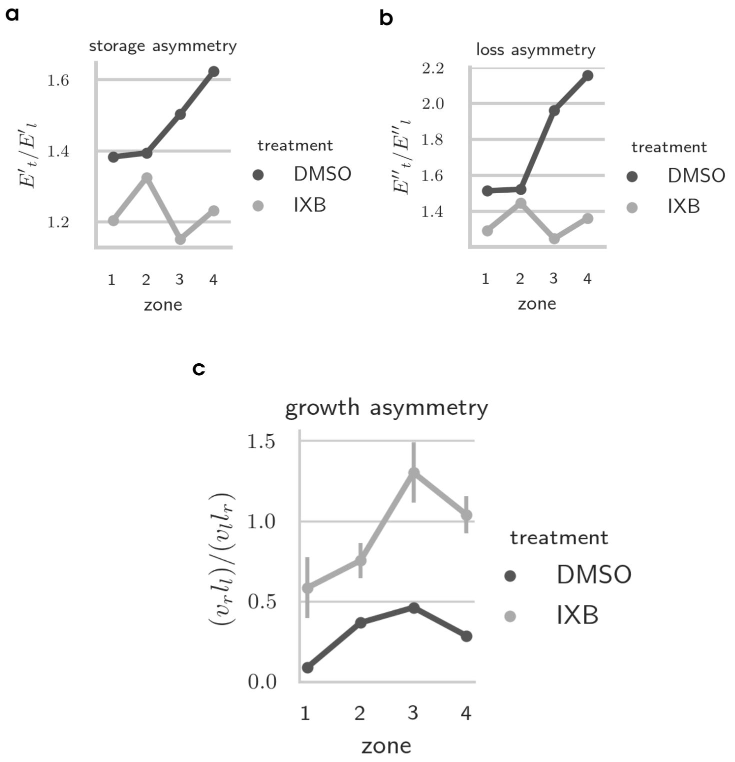


Figure S13: Asymmetry of IXB treated seedlings. Maps of storage (a) and loss (b) moduli as well as growth (c). The asymmetry is determined by the ratis of transverse to longitudinal walls for the moduli and radial to longitudinal relative growth rate for growth.

(11) M. Smith. *ABAQUS/Standard User's Manual, Version 6.9*. Dassault Systèmes Simulia Corp, United States, 2009.

(12) I. N. Sneddon. The relation between load and penetration in the axisymmetric boussinesq problem for a punch of arbitrary profile. *International Journal of Engineering Science*, 3(1):47–57, 1965. ISSN 00207225. doi: 10.1016/0020-7225(65)90019-4.
